# In vitro reconstitution of vertebrate Sonic Hedgehog protein cholesterolysis

**DOI:** 10.64898/2026.03.09.710561

**Authors:** Dayton C. Seidel, Andrew G. Wagner, John L. Pezzullo, Katherine A. Thayer, Seth Beadle, Margot L. Olejarczyk, José-Luis Giner, Brian P. Callahan

## Abstract

Extracellular secretion of the oncogenic sonic hedgehog signaling ligand is contingent on its release from a precursor protein through peptide bond cholesterolysis, mediated by the hedgehog C-terminal domain, SHhC. In this work, we describe the in vitro reconstitution of cholesterolysis activity for SHhC domains from vertebrate model organisms, *Xenopus laevis* (Xla) and *Danio rerio* (Dre). Cholesterolysis is assayed continuously in multi-well plates by monitoring changes in fluorescence resonance energy transfer (FRET) from an engineered precursor construct, expressed in *E. coli* and purified in soluble form. Using this FRET assay, we found that Xla and Dre SHhC exhibit high substrate stereospecificity, accepting cholesterol, (K_M_, 1-2 µM, cholesterolysis t_1/2_ of ∼11 min) while rejecting the 3-alpha epimer, epi-cholesterol (K_M_ > 100 µM, t_1/2_ > 10 hr). By screening a 96-member detergent/surfactant library for compatibility with SHhC activity, we identify cationic detergents that inhibit cholesterolysis and find a shared preference for the zwitterionic n-dodecyl-phosphocholine (DPC, Fos-choline-12), which supported the fastest reaction kinetics. Lastly, we report that alanine point mutation at a conserved aspartate residue (D46A) in Xla SHhC and Dre SHhC blocks cholesterolysis; however, activity could be chemically rescued with rationally designed hyper-nucleophilic sterols. Of those sterols, 2-beta carboxy cholestanol was active as a substrate with D46A variants only; the remaining sterols were accepted by both D46A and wild-type SHhC. In summary, we have established the first in vitro kinetic assay to continuously monitor enzymatic activity of wild-type and mutant vertebrate SHhC domains in multi-well plates, a key step toward pharmacological manipulation of Sonic hedgehog protein biosynthesis in vivo.

## INTRODUCTION

The extracellular Sonic hedgehog ligand (SHhN) initiates vital cell/cell signaling in pre-natal development through adulthood.^1^ Congenital mutations that diminish Sonic hedgehog protein expression are associated with holoprosencephaly, a potentially fatal brain disorder,^2, 3^ while chronic, overexpression appears to promote the growth of certain tumors._4-6_

Human SHhN and homologous proteins of metazoans are expressed as a bifunctional precursor with the hedgehog ligand flanked by a C-terminal cholesterolysis domain, SHhC. ^7–11^ Before SHhN ligand can be secreted extracellularly for biological signaling, SHhC must cleave itself from SHhN and covalently join the ligand’s newly formed carboxy terminus to a molecule of cholesterol (**Scheme 1**).^12, 13^ This unusual cleavage/sterylation event occurs site-specifically at a conserved Gly(-1)↓Cys(1) motif that separates SHhN from SHhC.^14, 15^ Residues are numbered here relative to SHhC, where the first amino acid of SHhC is 1 and the last residue of SHhN is −1.

As depicted above, cholesterolysis begins with isomerization of the Gly(-1)-Cys(1) backbone peptide bond, activating the Gly(-1) as an internal thioester. Following substrate cholesterol binding, the lipid’s 3-ϐ hydroxyl group is subjected to general base catalysis by a conserved aspartate residue (D46) of SHhC. Displacement of the internal thioester by cholesterol completes the chemical transformation.^1^ No cofactors/coenzymes are required. Cholesterolysis appears to be single turnover in the cell and entirely self-catalyzed by SHhC.^7, 14, 15^ The SHhN ligand adjacent to the cleavage site is a bystander; its replacement with unrelated polypeptides leaves SHhC activity intact in vitro and in vivo.^16–20^ Mutations that disable SHhC^21^ or extreme depletion of endogenous cholesterol,^22^ result in retention of unprocessed SHhN-SHhC in the secretory pathway where the precursor is targeted for degradation by the proteasome, blocking downstream signaling.^23, 24^

Biochemical studies, mutational analysis and small molecule discovery efforts on hedgehog cholesterolysis have focused on the *Drosophila melanogaster* domain, Dme HhC. ^17, 25–29^ Initial biochemical evidence for cholesterolysis was collected using this purified domain.^15^ Dme HhC is reliably overexpressed in *E. coli* in soluble, cholesterolysis active form and tolerates N- and C-terminal fusions to heterologous proteins and tag sequences.^17^ To date, Dme HhC remains the only cholesterolysis domain with experimentally determined high resolution structure data,^14, 30, 31^ although this information is limited to the protein’s first ∼150 amino acids; the final ∼80 amino acids of Dme HhC, which are necessary for recognition of substrate cholesterol, have so far resisted structural analysis.

*Drosophila* HhC and human SHhC share only 32% amino acid identity (45% similarity). Unlike Dme HhC, the native folding of human SHhC appears to require protein disulfide isomerase^23^ and asparagine glycosylation,^32^ features that may explain challenges in the heterologous expression of functional human SHhC in *E. coli*. It also seems worth noting that *Drosophila melanogaster* is a sterol auxotroph,^33^ a characteristic that could broaden Dme HhC substrate tolerance for (scavenged) sterols relative to human SHhC. Indeed, the native substrate for insect HhC domains is not known and may be a fungal or phytosterol.

Identifying SHhC cholesterolysis domains that are experimentally tractable and more closely related to human SHhC remains an important goal. In this work, we pursue SHhC domains from the vertebrate model organisms, *Xenopus laevis* (African clawed frog) and *Danio rerio* (Zebrafish). Pairwise comparisons of *Xenopus laevis* and *Danio rerio* SHhC domains with human SHhC show 49% identity/59% similarity and 46% identity/60% similarity, respectively (**Supporting Table 1**, **Supporting Figure 1**). Similar to human SHhN, the mature Zebrafish and Xenopus SHhN ligands are derived from SHhN-SHhC precursor cholesterolysis and regulate complex multicellular behavior. The zebrafish SHhN ligand participates in retinal development^34^, tooth formation^35^, and neurogenesis^36^. Signaling by *Xenopus laevis* SHhN is implicated in organogenesis^37, 38^ and tissue regeneration^39^.

**Figure 1.**
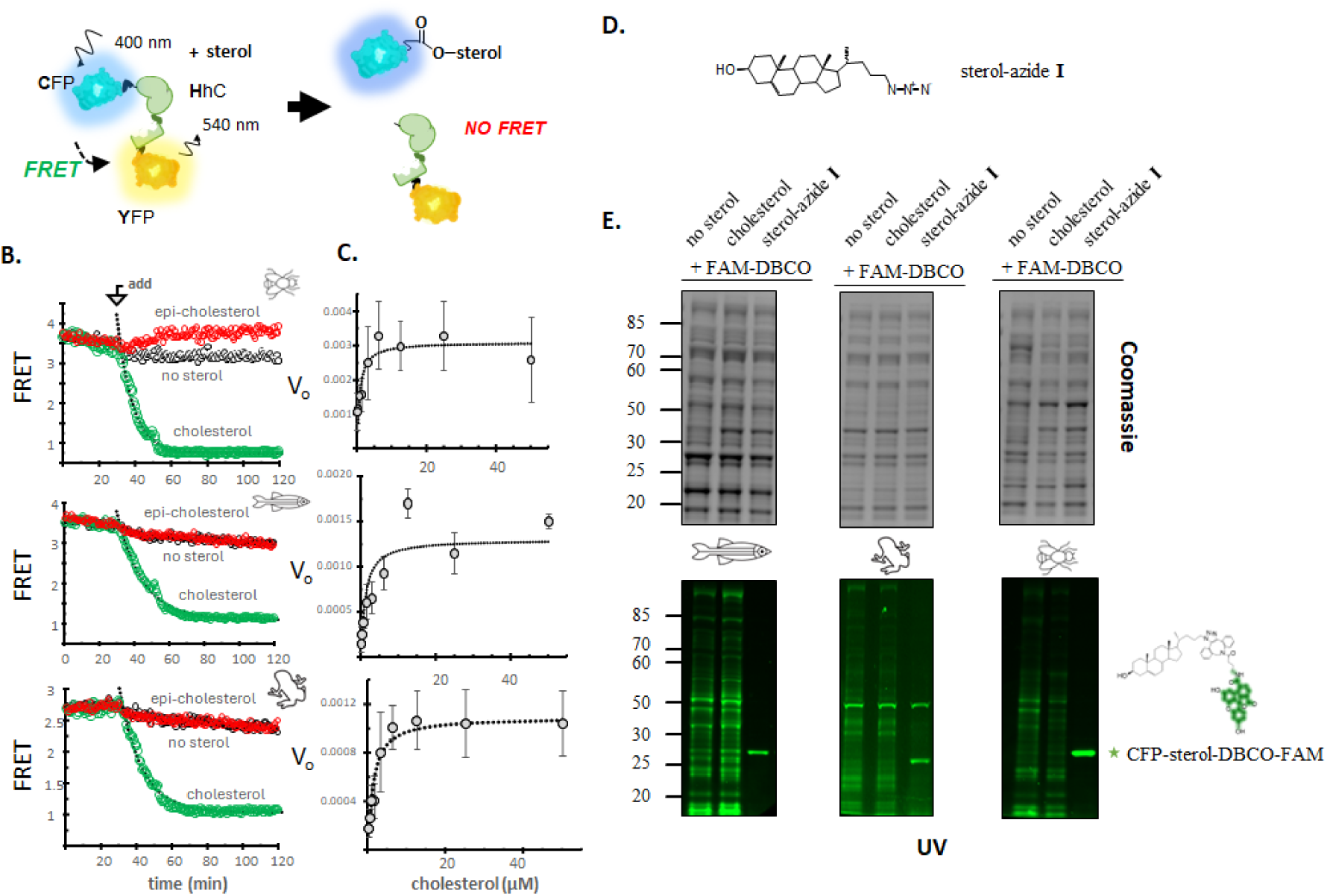
*Continuous in vitro assay of vertebrate SHhC cholesterolysis activity*. **A.** FRET-active cholesterolysis reporter. **B.** Kinetic assays with substrate cholesterol and inactive epi-cholesterol. After a thirty-minute incubation of samples containing C-H-Y reporter, sterol was added from an ethanol stock to final concentration of 50 µM, 2% ethanol. “No sterol” control reactions had the same volume of ethanol added to evaluate possible solvent effects. Representative kinetic traces are shown for wild-type C-H-Y reporters for *Drosophila melanogaster* (Dme), *Danio rerio* (Dre) and *Xenopus laevis* (Xla). **C.** Concentration-response plots for cholesterolysis. Wild-type C-H-Y reporter constructs with Dme, Dre and Xla domains exhibited Michaelis-Menten type behavior in plots of initial rate of FRET-loss as a function of increasing initial cholesterol concentration. Derived K_M_ values were ∼ 1 µM. **D**. Structure of clickable cholesterol azide analog. **E**. Covalent sterylation. Clickable cholesterol (I) was used as an alternative substrate for wild-type C-H-Y reporter constructs of Dme, Dre and Xla in the presence of E. coli soluble lysate. Completed reactions were mixed with DBCO-FAM, 1:1 with (**I**), separated by denaturing SDS-PAGE and imaged for fluorescence using BioRad EZ Gel Doc. Negative control reactions without (I), showed an unspecific laddering of the fluorescent label (lanes 1-2). Samples with clickable sterol showed a prominent band at the expected molecular weight for sterylated CFP (lane 3).

**Table 1.** Cholesterol substrate activity with wild-type SHhC domains at 30 °C, pH 7.1 using Fos-choline-12 (1.5 mM) as detergent.

| | $k_{\text{obs}}$ (s<sup>-1</sup>) | $K_M$ (μM) |
| --- | --- | --- |
| *Dme HhC* | 1.2*10<sup>-3</sup> | 0.69 |
| *Dre SHhC* | 1.1*10<sup>-3</sup> | 1.1 |
| *Xla SHhC* | 1.1*10<sup>-3</sup> | 1.3 |

As a step toward pharmacological manipulation of sonic hedgehog protein biogenesis, we establish here the first continuous, in vitro kinetic assays to monitor *Xenopus laevis* and *Danio rerio* SHhC cholesterolysis. Assays are conducted in multi-well plates with an engineered FRET-active SHhC reporter produced in soluble form using *E. coli*. Compared to conventional gel-based SHhC activity assays, the FRET-method improves the speed of sample throughput and provides a more complete picture of reaction progress through real-time data acquisition. We use this method to show that vertebrate SHhC domains, like their invertebrate counterpart Dme HhC, exhibit low micromolar affinity for substrate cholesterol while rejecting epi-cholesterol, the 3-epimer of cholesterol. We observed that both vertebrate SHhC domains are susceptible to precursor thiolysis in the absence of cholesterol, suggesting that internal thioester formation and substrate binding by SHhC are not strictly coupled. Screening a 96-member library of potential sterol solubilizing agents for in vitro SHhC cholesterolysis, we found that vertebrate and invertebrate cholesterolysis domains displayed a shared preference for the phosphatidylcholine analogue, fos-choline 12 (DPC). Lastly, we report that alanine point mutation at a conserved aspartate residue (D46) in the zebrafish and Xenopus SHhC domains allows for bioorthogonal “chemical rescue”, where SHhC cleavage/sterylation activity is dependent on exogenous hyper-nucleophilic sterol.

## RESULTS AND DISCUSSION

### Heterologous *E. coli* expression of vertebrate SHhC reporter constructs

To explore the viability of *in vitro* enzymatic studies involving *Xenopus laevis* and *Danio rerio* SHhC domains, hereafter Xla SHhC and Dre SHhC, we cloned codon optimized fragments encompassing the respective proteins along with a short flanking SHhN peptide into an arabinose-inducible FRET reporter expression plasmid, extending the approach we used previously for *Drosophila melanogaster* HhC (Dme HhC).^17^ The encoded polyprotein, C-H-Y, has an N-terminal cyan fluorescent protein (C), followed by the hedgehog SHhC domain (H), fused to yellow fluorescent protein (Y) with a C-terminal His6 purification tag (**Supporting Figure 2**).

**Figure 2.**
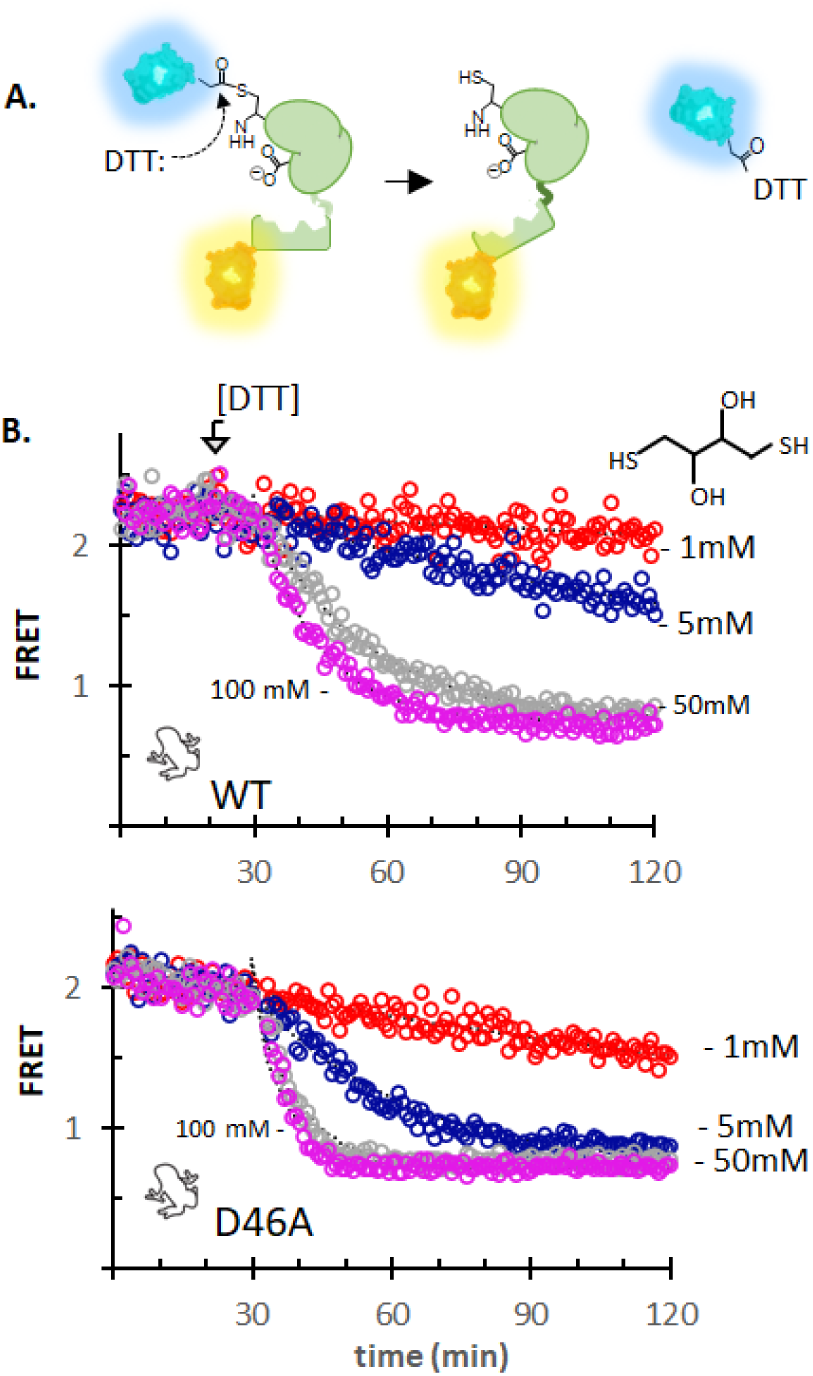
*Sterol-independent thiolysis of WT and D46A SHhC precursor*. **A**. Dithiothreitol (DTT) cleavage of FRET-active C-H-Y precursor into C and H-Y fragments with ensuing loss of FRET. **B**. Kinetic traces for cleavage of Xla C-H-Y wild-type (top) and Xla C-H-Y D46A (bottom) with increasing concentration of DTT. Reactions were carried out at 30 °C under identical conditions to cholesterolysis assays of Figure 1 except that cholesterol was absent.

Along with wild-type FRET reporter constructs, we prepared two alanine point mutants of Xla SHhC and Dre SHhC: a C1A mutant to serve as a negative control where the catalytically essential nucleophilic thiol of SHhC is removed; and a D46A mutant, where the putative general base function of the D46 side-chain carboxyl group is eliminated. The D46A mutation in *Drosophila* HhC and human SHhC have proven amenable to chemical rescue of their cleavage/sterylation activity.^16, 40^

FRET reporter constructs of wild-type and the two point mutants of Xla and Dre SHhC were overexpressed in *E. coli* and purified in soluble form using standard Ni-NTA chromatography (**Supporting Figure 3**). Compared with Dme HhC, yields of purified C-H-Y incorporating the vertebrate domains were reduced by a factor of 4.5 and 3 for Xla SHhC and Dre SHhC respectively. Two different E. coli strains were tested, LMG194 and AI-BL21; yields were slightly better in AI-BL21. Analysis of elution fractions by SDS-PAGE indicated that spontaneous, hydrolytic autoprocessing of the Xla and Dre SHhC constructs was greater than with Dme HhC. Despite reduced yields with the vertebrate domains, the WT, C1A and D46A SHhC constructs exhibited FRET signal five to seven-fold above background and displayed the expected cholesterolysis activity, inactivity, or conditional activity, as described below.

**Figure 3.**
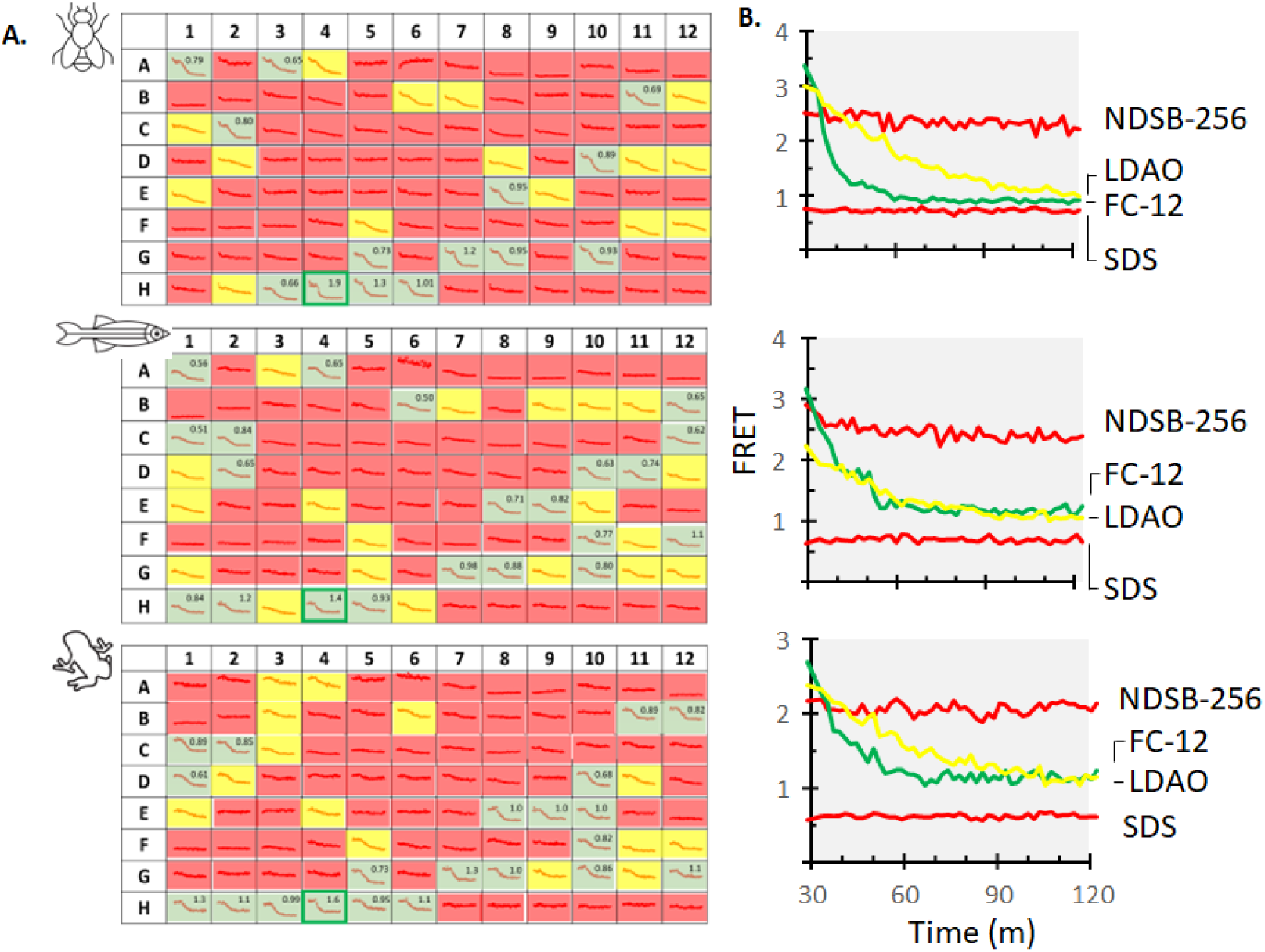
*Detergent selectivity is similar for invertebrate and vertebrate cholesterolysis domains*. **A**. Cholesterolysis activity for Dme, Dre and Xla domains using the C-H-Y reporter in the presence of 96 potential sterol solubilizing agents. Each detergent/surfactant was added at their critical micelle concentration. Kinetics were scored qualitatively as: fast (green); moderate (yellow); weak or no activity (red). First order rate constants (x10^-3^ sec^-1^) are shown in the upper right corner for “green” samples. The detergent fos-choline 12 supported the fastest kinetics across the three proteins (well, H4). **B**. Representative kinetic traces for green, yellow, and red detergents. Plots ordered as followed: Dme (top), Dre (middle) and Xla (bottom). Abbreviations: NDSB-256 (non-detergent sulfobetaine); LDAO (N,N-dimethyl-1-dodecanamine, N-oxide); FC-12 (n-Dodecylphosphocholine; Fos-choline-12); SDS (Sodium Dodecyl Sulfate).

### Vertebrate hedgehog cholesterolysis activity in vitro by continuous FRET assay

Xla and Dre SHhC cholesterolysis have been examined before using label-based, endpoint approaches. Xla SHhC activity was detected in the presence of Xenopus egg extracts with in vitro translated ^35^S-labeled Xenopus Sonic Hh precursor protein.^23^ A nonradioactive chemical tagging approach involving an alkynyl-modified cholesterol, click chemistry and in-gel fluorescence, was developed for Dre SHhC cholesterolysis. ^41^

For higher throughput and continuous monitoring of reaction progress, we sought to assay Xla and Dre SHhC cholesterolysis in multi-well plates using a fully recombinant, label-free reporter protein. The schematic in **Figure 1A** depicts the central reagent C-H-Y and its operation. In the unprocessed C-H-Y precursor, the hedgehog cholesterolysis domain (H) maintains the FRET donor, CFP, and the FRET acceptor, YFP, in proximity, resulting in a stable FRET signal. We follow the convention of reporting FRET as an emission ratio of 540 nm/460 nm after 400 nm excitation.^42^ If the “H” domain of C-H-Y is functional, the FRET signal is expected to decay in a time dependent manner following the addition of substrate (X), reporting the separation of product CFP-X from H-Y. Assays are conducted at 30 °C in 96-well plates with C-H-Y (0.2-0.1 µM) in Bis–Tris buffer (pH 7.1) and Fos-choline 12 (1.5 mM, final), the latter added as a sterol solubilizing agent. To suppress cysteine oxidation, EDTA (5 mM, final) and TCEP or DTT (4 mM, final) are included. We routinely incorporated a 10 to 30 min preincubation period with C-H-Y in complete assay buffer minus substrate to establish a starting FRET value. Following the addition of substrate, usually from an ethanol stock solution, FRET readings are recorded every 1 to 2 minutes.

As shown in **Figure 1B**, we observed stereospecific cholesterolysis activity for wild-type Xla and Dre SHhC using the FRET system. *Drosophila* Dme HhC in the C-H-Y construct was included as a control and comparator in all experiments. Stable FRET signal from samples in the preincubation period without cholesterol is consistent with successful folding of the C, H, and Y elements of the precursor protein. Cleavage of C-H-Y was apparent following the addition of excess cholesterol, 250x > [C-H-Y], which resulted in time dependent loss of FRET signal that followed pseudo-first order kinetics (**Figure 1B**, green symbols). The derived rate constants, k_max_, were as follows: 0.0012 sec^-1^ for *Drosophila*, and 0.0011 sec^-1^ for the zebrafish and *Xenopus* domains. Solvent control reactions where an equivalent volume of ethanol without cholesterol was added to C-H-Y samples did not appreciably alter FRET signal over the time course of the experiment (**Figure 1B**, black symbols). Similar inactivity was apparent in C-H-Y samples mixed with epi-cholesterol (**Figure 1B**, red symbols).^12^ The C1A mutants of Xla and Dre SHhC were not reactive with cholesterol or epi-cholesterol, consistent with the imperative for an N-S acyl shift to activate the G(-1) residue for cleavage (**Supporting Figure 4**). The D46A mutants of Xla and Dre SHhC also appeared unreactive with cholesterol up to 100 µM.

**Figure 4.**
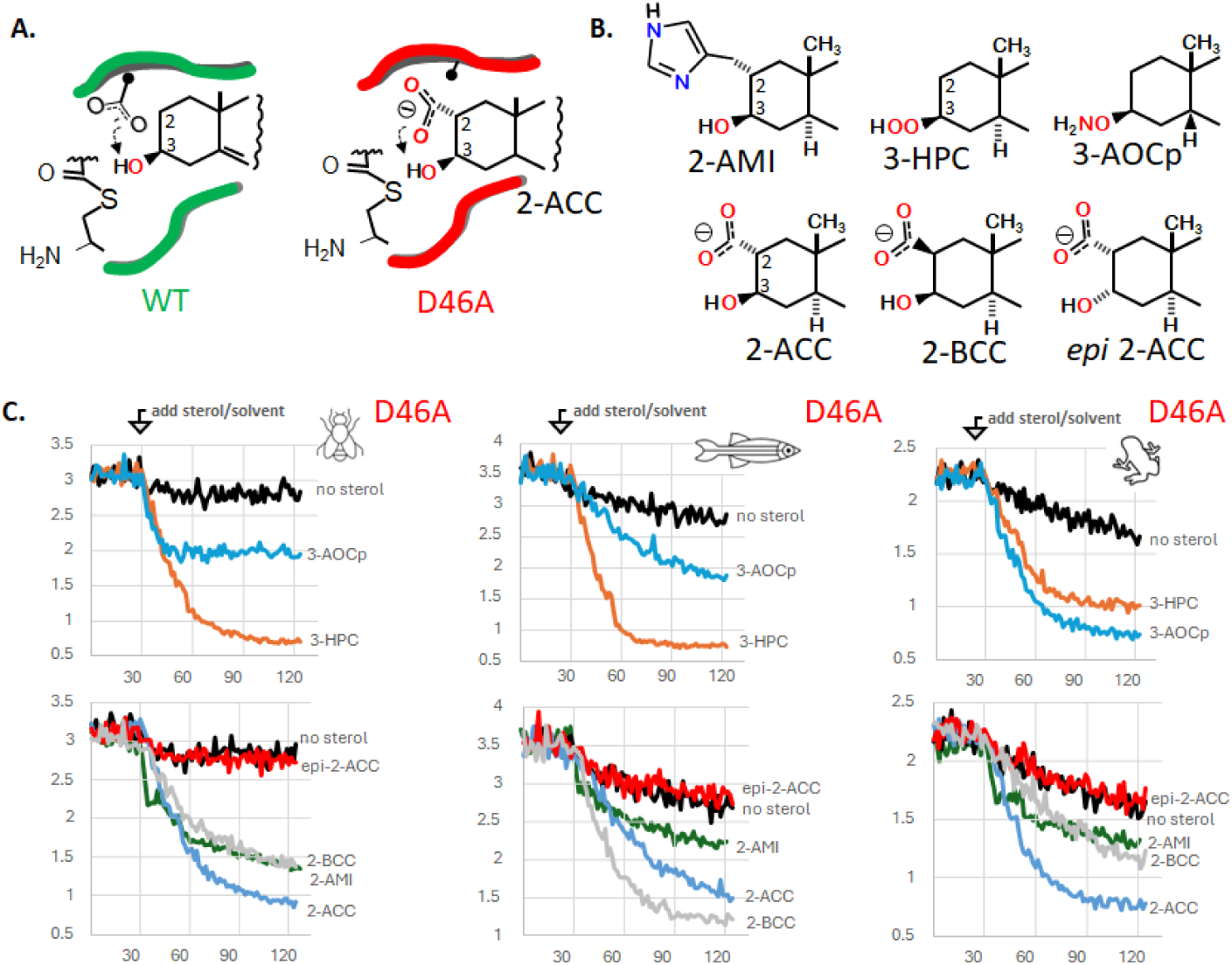
*Chemical rescue of D46A mutants using hyper-nucleophilic sterols*. **A**. Putative catalytic base role of conserved D46 (left); bypass pathway in D46A mutant using synthetic carboxy sterol, 2-alpha carboxy cholestanol. **B**. Chemical rescue and negative control sterols. Abbreviations: 2-AMI, 2-alpha methyl imidazole cholestanol; 3-HPC, 3-beta hydroperoxyl cholestanol; 3-AOCp, 3-beta aminoxy coprostanol; 2-ACC, 2-alpha carboxy, 3-beta hydroxy cholestanol; 2-BCC, 2-beta carboxyl, 3-beta hydroxy cholestanol; epi-2ACC, 2-alpha carboxy, 3-alpha hydroxyl cholestanol. **C**. Representative kinetic traces with D46A C-H-Y reporters for Dme, Dre and Xla domains with indicated sterol. (Top) Rescue sterols bearing alpha effect nucleophiles at the 3-position. (Bottom) Rescue sterols bearing potential general base catalysts at the 2-position. After a thirty-minute preincubation, sterols were added to a final concentration of 50 µM. Fos-choline-12 was used as the solubilizing agent.

The derived rate constant for Xla SHhC cholesterolysis is in reasonable agreement with the kinetics we estimate from the study with ^35^S labeled Xla SHh precursor in egg extract. ^23^ For additional comparison, an apparent rate constant of 0.0005 sec^-1^ for Human SHhC cholesterolysis was reported from cellular pulse-chase experiments,^22^ which is within 3-fold of the values reported here.

Concentration-response plots of the initial rate of FRET loss with increasing cholesterol showed saturation behavior (**Figure 1C**). Derived K_M_ values of Xla SHhC and Dre SHhC were 1.3 µM and 1.1 µM, comparable to the cholesterol K_M_ value for Dme HhC (**Table 1**). In other enzyme systems, the K_M_ value can approximate the substrate concentration in vivo.^43^ The degree of substrate saturation for hedgehog cholesterolysis in the secretory pathway is not known. In earlier work, we observed EC50 values of ∼6 µM for exogenous sterol added to chemically rescue a conditional mutant of human sonic hedgehog SHhC (D46A) expressed in HEK293 cells.^16^ A similar concentration of exogenous sterol analogs was used in mammalian cell experiments as alternative substrates for SHhC cholesterolysis.^44^ In summary, the cholesterol K_M_ values from the in vitro reconstituted SHhC assays are in the low micromolar range, in general agreement with prior cellular studies.

To validate covalent sterylation of the product “C” protein from vertebrate SHhC containing C-H-Y precursors, we used 24-azide-modified cholesterol analog (I) ^45–47^ as substrate, followed by strain-promoted click chemistry with DBCO-fluorescein (**Figure 1D**). Resulting protein-sterol conjugates were then analyzed by denaturing SDS-PAGE followed by in-gel fluorescence. We carried out the sterolysis and the subsequent click reaction in total soluble E. coli lysate. In-gel fluorescence of control reactions with C-H-Y that lacked sterol, or used native cholesterol, showed a non-specific pattern of fluorescein labeling (**Figure 1E**, *first two lanes*).^2^ By contrast, a strong, distinct fluorescent protein band was apparent in samples containing C-H-Y, the azide-sterol and DBCO-fluorescein (**Figure 1E**, *third lane*). The position of the fluorescent protein bands in the three gels corresponds to the “C” protein molecular weight (∼30 kDa).

Taken together, results from this section indicate that recombinant SHhC domains of *Xenopus laevis* and *Danio rerio* can be expressed in *E. coli* in soluble, cholesterolysis-active form. Both vertebrate SHhC domains tolerate N- and C-terminal fusion to fluorescent proteins, enabling continuous FRET-based reporting of cholesterolysis kinetics in a multi-well format. This cleavage/sterylation activity of Xla and Dre SHhC domains with cholesterol requires their Cys1 and D46 residues, and the 3-OH group of the cholestane must have a ϐ configuration for recognition as a substrate.

### Thioester formation by SHhC does not require cholesterol binding

In vitro studies of purified Drosophila Dme HhC suggested that internal thioester formation (step 1, Scheme 1) is not dependent on the presence of sterol substrate.^14^ De-acylating agents such as hydroxylamine and dithiothreitol (DTT), for example, can site-specifically cleave purified Hh precursor protein at the Gly(-1)-Cys(1) junction in sterol-free buffer (**Figure 2A**).^14, 48^ Although much slower than thiolysis and hydroxyaminolysis, spontaneous hydrolytic cleavage at the same Gly(-1)-Cys(1) motif has also been observed (**Supporting Table 2**).^17, 27^ These non-native cleavage pathways of Dme HhC are eliminated by Cys1Ala mutation. It seems reasonable to infer from these results that the Hh precursor Gly(-1)-Cys(1) bond is an equilibrium mixture of backbone amide and side chain thioester irrespective of sterol substrate.

We found that Dre SHhC and Xla SHhC precursors were likewise susceptible to DTT-induced thiolysis in the absence of cholesterol. The three wild-type C-H-Y constructs were cleaved in C and H-Y fragments by 100, 50 and 5 mM DTT in a concentration dependent manner (**Supporting Figure 5**). **Figure 2** shows the DTT cleavage traces with Xla ShhC, WT and D46A. Zebrafish Dre SHhC reacted most rapidly with DTT, followed by Xla SHhC and Dme HhC. Dre SHhC displayed weak but measurable cleavage with 1 mM DTT, while Xla SHhC and Dme HhC were insensitive at this low DTT concentration.

**Figure 5.**
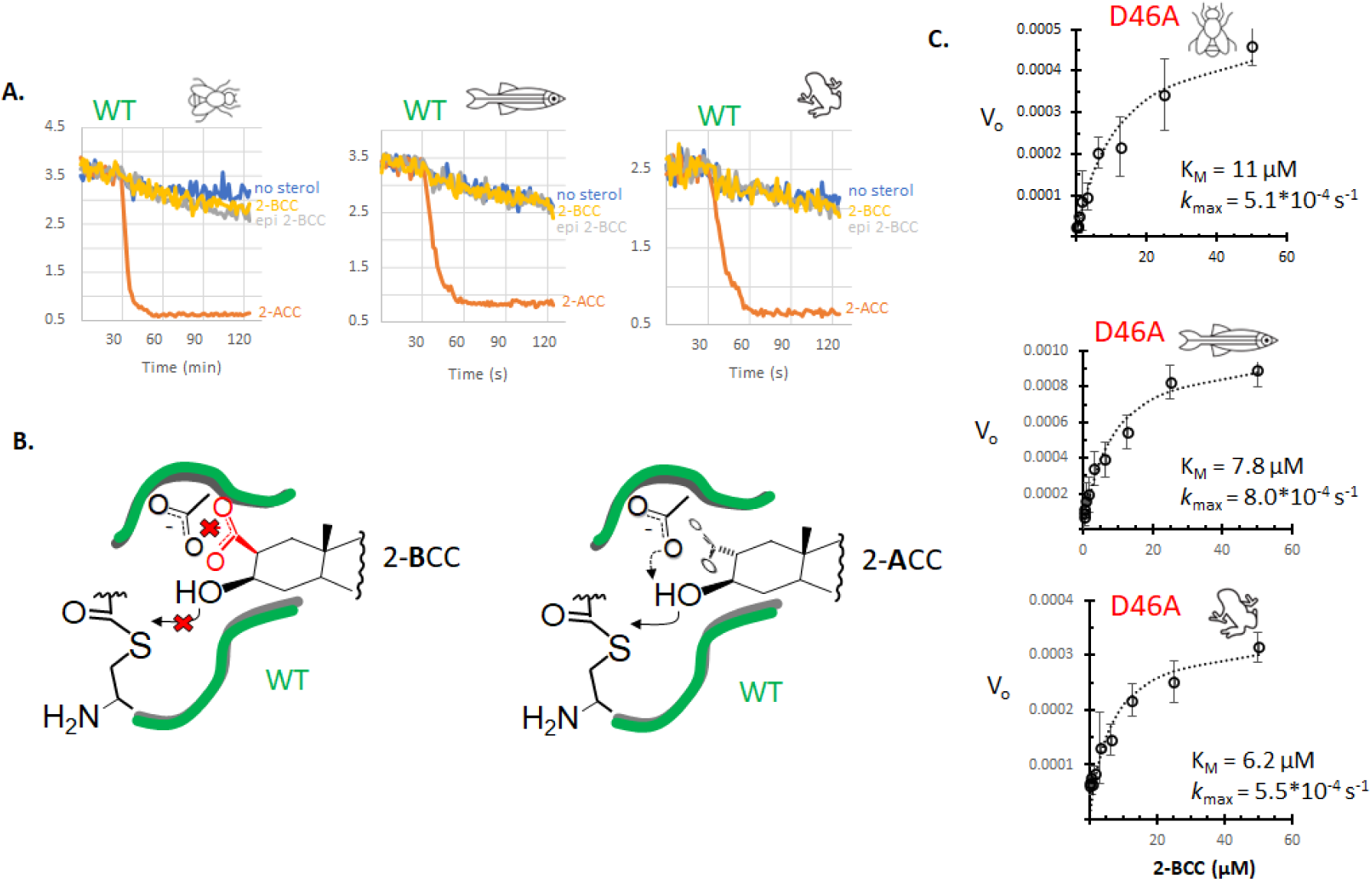
*D46A-specific chemical rescue with 2-BCC*. **A**. Wild-type cholesterolysis domains reject 2-BCC as a substrate. Dme, Dre and Xla C-H-Y reporter displayed robust substrate activity toward 2-ACC but not 2-BCC, or epi-2-BCC. Sterols were tested at 50 µM with Fos-choline-12 (1.5 mM, final). **B**. Proposed carboxyl-carboxyl repulsion that blocks 2-BCC but not 2-ACC from productive binding with wild-type cholesterolysis domain. **C**. Concentration response plots for 2-BCC substrate activity with invertebrate and vertebrate D46A SHhC domains along with their derived K_M_ and *k_max_* values.

Alanine substitution at D46 accelerated DTT-induced thiolysis by 2-6 fold in Dre and Xla SHhC variants. Previous structure-function studies with Dme HhC led to the proposal that the carboxylate side chain of D46 restricts N-to-S acyl shift activity at the Gly(-1)-Cys(1) motif in the absence of cholesterol. The net effect is to reduce the risk of wasteful hydrolytic cleavage of the internal Cys(1) thioester.^29^ Enhanced DTT-cleavage rates with D46A mutants of Dre and Xla SHhC suggests that this conserved residue plays a similar regulatory role for the vertebrate domains.

As expected, alanine mutation at thioester-forming Cys1 abolished DTT-induced cleavage of Dre SHhC and Xla SHhC (**Supporting Figure 4**). In summary, the cholesterol-independent precursor thiolysis results (**Supporting Table 3**) indicate that thioester formation at the Gly(-1)-Cys(1) bond and substrate binding by SHhC are not strictly coupled.

### Detergent compatibility / incompatibility for in vitro cholesterolysis

Cholesterol is sparingly soluble in aqueous solution (∼1 nM), ^49^ requiring the addition of a detergent to prevent its precipitation. In the SHhC assays above, Fos-Choline 12 was used as the sterol solubilizing agent, a selection based on the results of a preliminary screen for compatible detergents we carried out using Dme HhC.

To explore the extent to which detergent structure impacts SHhC cholesterolysis activity and with a view toward future structural biology and protein biotechnology applications, we screened a commercial 96-member detergent library for SHhC compatibility. The library includes 46 non-ionic, 9 anionic, 4 cationic, and 31 zwitterionic agents, as well as six non-detergent sulfobetaines (**Supporting Figure 6**). Each detergent was tested in the FRET assay at their critical micelle concentration.

Reaction progress curves for cholesterolysis were scored initially by visual inspection: *green*, rapid cholesterolysis; *yellow*, cholesterolysis active but moderate rate; *red*, no apparent cholesterol-dependent reaction (**Figure 3**). Some *Red* detergents, such as sodium dodecyl sulphate and sodium cholate, appeared to destabilize C-H-Y, as suggested by FRET values that started out low (<1.5) or trended lower during the assay preincubation period when cholesterol was absent. Following this qualitative assessment, kinetic data for all *green* detergents were analyzed quantitatively to obtain apparent first-order rate constant for SHhC cholesterolysis (**Supporting Table 4**).

Comparison of the results across the three homologous cholesterolysis domains showed similar behavior overall. Most detergents in the library were categorized as either *yellow* or *red* for weak or no activity. None of the cationic detergents were classified as *green* for high activity, and at least one (cetylpyridinium chloride, *well* **A5**) appeared inhibitory. The sulfobetaine molecules also failed to support cholesterolysis (*wells* **H7-H12**). The list of *yellow* detergents included Triton-X 100, which was used as a sterol solubilizing agent in early studies of Dme HhC cholesterolysis (*well* **E6**). Non-ionic detergents that supported more rapid cholesterolysis included nonylphenyl PEG for Dme HhC, and Tween 80 for Zebrafish and Xenopus SHhC (*wells* **E8** and **E10**). The non-ionic detergent, n-decanoyl D-sucrose, considered readily removable by dialysis, supported rapid kinetics for all three homologues (*well* **D10**). Among anionic detergents, 14:0 Lyso PG appeared best for Dme HhC, while the vertebrate Dre SHhC and Xla SHhC preferred 18:1 Lyso PG (*wells* **A1** and **A4**).

Fos-choline 12 (dodecylphosphocholine, or DPC), the synthetic zwitterionic detergent (*well* **H4**), supported the fastest kinetics for Dre SHhC, Xla SHhC and Dme HhC. In the ER, where SHhC encounters substrate cholesterol, structurally analogous phospholipids are abundant, providing a possible biological rationale for this selectivity. The effect of Fos-choline 12 may involve an activating conformational change of SHhC. Preliminary concentration-response plots for C-H-Y, where FRET is plotted as a function of increasing Fos-choline 12 concentration, showed increasing FRET signal that peaks around the CMC value for Fos-choline 12 (**Supporting Figure 7**). A control construct, C-Y, where cyan and yellow fluorescent proteins are fused without SHhC, lacked a comparable trend in FRET signal with increasing Fos-choline 12. The molecular details of the Fos-choline 12 and SHhC interaction awaits more detailed structural analysis. To that end, Fos-choline 12 is often used for solution NMR of membrane binding proteins. Other zwitterionic detergents that scored *green* for fast kinetics with Dre SHhC, Xla SHhC and Dme HhC included 12:0 Lyso PC, 13:0 Lyso PC, and 15:0 Lyso PC, as well as Fos-choline 14 (*wells* **G7**, **G8**, **G10**, and **H5**).

To summarize, detergent structure can have a profound effect on the rate of SHhC cholesterolysis. Zwitterionic detergents appear to be the favored charge type for the invertebrate and vertebrate domains, while the cationic detergents tested here lacked SHhC compatibility. Although the overall trends are similar, we did identify species specific preferences with certain detergents (**Supporting Figure 8**). For in vitro assays of SHhC cholesterolysis, the zwitterionic Fos-choline 12 is recommended.

### Chemical rescue of SHhC D46A mutants with hyper-nucleophilic sterols

Chemical rescue is a mechanism-guided approach to probe enzymatic activity. A catalytically essential functional group is removed first by alanine point mutation, compromising activity toward the native substrate. Conventional chemical rescue strategies graft the missing functional group onto the substrate molecule^50^ or add the functional group in *trans* as a makeshift co-factor. ^51^ In an early example of the latter approach, a catalytic mutant of aspartate aminotransferase where alanine replaced the key general base, K258, enzymatic activity was chemically rescued by addition of exogenous aliphatic amines. ^50^ Additional examples have been reviewed.^52^ We recently introduced a third strategy for chemical rescue that uses “alpha effect” nucleophiles,^40^ which we apply again here.

The carboxylate group of SHhC’s D46 residue is essential for native cholesterolysis. As shown in **Figure 4A**, the carboxylate side chain supplies a general base to deprotonate the substrate’s 3-beta hydroxyl group. Accordingly, D46A mutants of Dme HhC and human SHhC lack measurable activity toward cholesterol. Similar inactivity was observed after D46A substitutions in Dre SHhC and Xla SHhC. As apparent from the rapid rates of precursor thiolysis by DTT (**Figure 2**), however, D46A variants retain their ability to form internal thioester. In two earlier studies, the first involving Dme HhC (D46A) and more recently with human SHhC(D46A) expressed in HEK293 cells, we reported that cleavage/sterylation activity could be chemically rescued by substituting cholesterol with 3-hydroperoxycholestanol (3-HPC) and 2-alpha carboxyl cholestanol(2-ACC), respectively. ^16, 40^ These nonnatural hyper-nucleophilic sterols were designed to bypass the missing general base function of D46.

Chemical rescue is extended here to Xla and Dre SHhC D46A variants using 2-ACC, 3-HPC and three new rescue sterols (**Figure 4B**). 2-alpha carboxyl cholestanol (2-ACC) was designed for intra-molecular general base catalysis of the sterol’s 3-β hydroxyl group. To further test this chemical rescue mechanism, we prepared 2-alpha methyl imidazole cholestanol (2-AMI) and 2-beta carboxy cholestanol (2-BCC). Similar to 2-ACC, the synthetic sterols restored activity to D46A variants of Xla and Dre SHhC as well as Dme HhC. Chemical rescue is stereospecific, as the 3-epimer of 2-ACC was inactive as a substrate. Next, we tested 3-HPC and a related analogue, 3-AOCp. The hydroperoxyl group in 3-hydroperoxycholestanol (3-HPC) is an α-effect nucleophile, where the attacking atom is adjacent (i.e., alpha) to a second electronegative atom. In a similar way, the aminoxy group of 3-aminoxycoprostanol is an α-effect nucleophile. Hydroperoxy and aminoxy groups are recognized for reacting at exceptionally fast rates compared to non-α-effect nucleophiles of equal basicity.^53–55^ We found that both 3-HPC and 3-AOCp restored activity to D46A mutants, encouraging further exploration of α-effect chemical rescue.

Conditional SHhC mutants hold promise as tools to control hedgehog cell/cell signaling in vivo. As mentioned earlier, extracellular release of the sonic hedgehog signaling ligand (SHhN) is contingent on cleavage from and sterylation by SHhC. If cholesterol and other endogenous sterols fail to serve as substrates, as with D46A mutants, the SHhN ligand is retained intracellularly and degraded. Addition of synthetic sterols that can chemically rescue D46A cleavage/sterylation activity might restore SHhN ligand release and downstream hedgehog signaling. A potential complication for this longer-term goal arises from the observed activity of 2-ACC, 2-AMI, 3-HPC, and 3-AOCp with both D46A and wild-type SHhC (**Supporting Figure 9-10**, **Supporting table 5**). In vertebrate organisms like *Xenopus laevis* and *Danio rerio* that express multiple Hh proteins, addition of non-natural sterols that promiscuously activate WT and D46A SHhC might produce broad and potentially undesired, hedgehog pathway activation. We observed a notable exception to this promiscuity with 2-BCC.

### Orthogonality of Chemical rescue with 2-beta carboxy cholestanol (2-BCC)

2-β carboxyl cholestanol showed > 100-fold selectivity for SHhC(D46A) over the wild-type protein, suggesting the possibility of truly orthogonal, mutant-specific chemical rescue. The lack of wild-type SHhC activity toward 2-BCC (**Figure 5A**) may arise from charge-charge repulsion between the carboxylate groups of the native D46 side chain and 2-BCC, thereby interfering with productive substrate interactions (**Figure 5B**). ^3^ This proposed unfavorable interaction would be absent in D46A variants. Plots of initial reaction velocity using 2-BCC as a substrate for D46A variants are shown in **Figure 5C**. For all three proteins, we found K_M_ and k_max_ values that were within a factor of ten of the corresponding values for cholesterol with the WT constructs. The potential for the 2-BCC/D46A pairing to manipulate sonic hedgehog ligand biogenesis and downstream signaling in complex living systems is being examined.

## CONCLUSION

Peptide-bond cholesterolysis is a defining feature of sonic hedgehog ligand biosynthesis. While extracellular leakage of precursor hedgehog (SHhN-SHhC) has been observed in cell-culture studies, the strong genotype/phenotype association of SHhC deactivating mutations with human holoprosencephaly indicates that unprocessed hedgehog protein is inadequate for full biological signaling in vivo.^2, 3, 21, 24^ The C-terminal cholesterol molecule attached to SHhN by SHhC provides an important recognition element for extracellular SHhN transporters, assists in forming morphogenic gradients of hedgehog signaling, along with supporting functional interactions with SHhN cell surface receptors. ^56–62^

Small molecule activators of SHhC that can boost SHhN biosynthesis garner interest for mitigating hedgehog mutations linked to holoprosencephaly, whereas inhibitors of SHhC cholesterolysis activity hold promise for tamping down oncogenic SHhN biosynthesis and signaling.^16^ To date, no specific agonist or antagonist of human SHhC has been reported.

As the hedgehog founding family member, the *Drosophila melanogaster* HhC domain has long served as a valuable surrogate for understanding the molecular mechanisms of human SHhC activity. Nevertheless, sequence divergence between the human and fly proteins has motivated the pursuit of experimentally tractable SHhC domains from closer human homologues.

In this report, we established the first continuous in vitro cholesterolysis assays for SHhC domains from two vertebrate model organisms, *Danio rerio* and *Xenopus laevis*. The central reagent is a label-free, recombinant FRET-active reporter construct that monitors SHhC activity in real time, replacing more cumbersome gel-based endpoint assays. With this approach, we carried out initial kinetic characterization of vertebrate SHhC cholesterolysis and thiolysis activity, identified compatible detergents for SHhC activity, and extended the panel of unnatural sterols for chemical rescue of SHhC (D46A) mutants. Of special importance is the behavior of 2-ϐ carboxyl cholestanol, representing the first mutant-selective substrate for hedgehog precursor cleavage/sterylation.

## Acknowledgements

We are grateful for technical support and encouragement from past and present members of the Callahan lab, particularly Adam Elfadel and Dr. Dan Ciulla. This work was supported by NCI grant 5R21CA282857 (B.P.C.); J.-L.G. gratefully acknowledges support from the NIH Grant R15GM143714 and NIH S10 OD012254 for the 800 MHz NMR spectrometer.

## Footnotes

1 Recently, Wang and colleagues reported experimental evidence for an additional chemical step, preceding displacement by cholesterol, where the C(1) thioester migrates to a second conserved cysteine residue of SHhC to form a “branched intermediate”. REF 26

2 Addition of azido-PEG1 (50 µM, final) to control reactions lacking the sterol-azide probe (lanes 1, 2), eliminated this unspecific fluorescence labeling.

3 Consistent with anion-anion charge repulsion, initial characterization of an analog of 2-BCC where the imidazolyl analog is accepted by both D46A and WT proteins, with enhanced reactivity toward WT

---

**Scheme 1.**
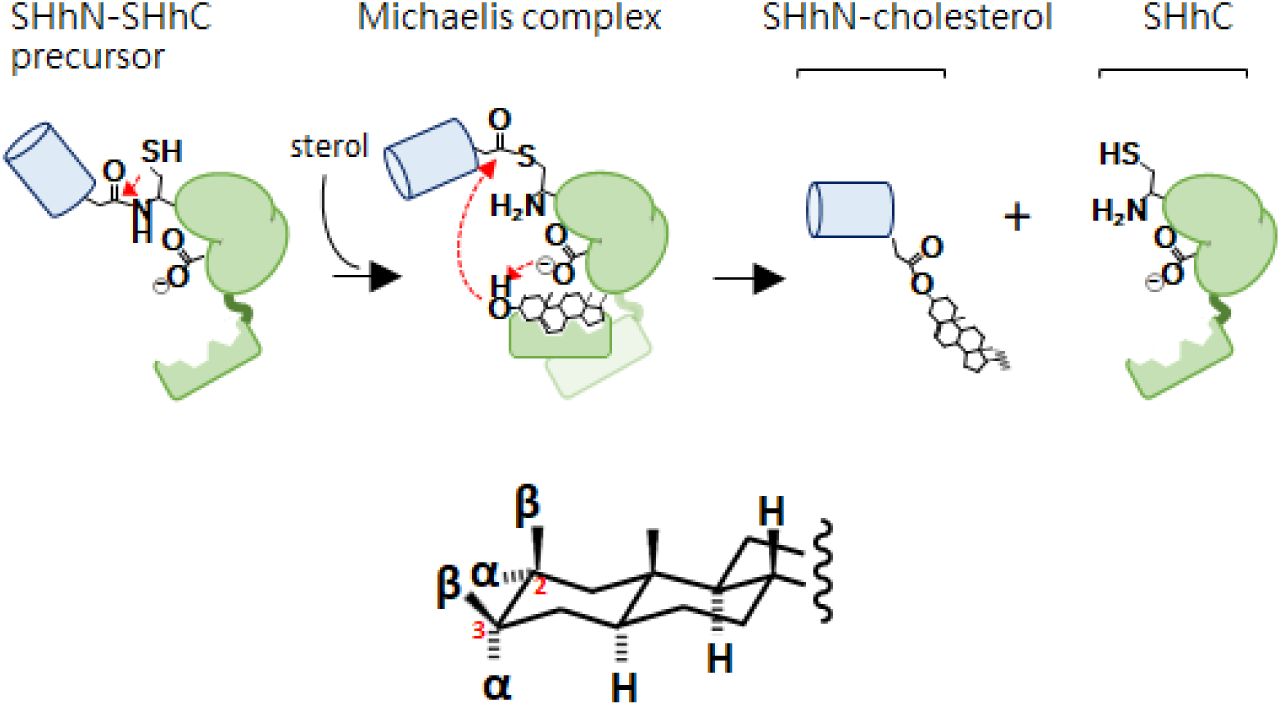
(top) Sonic hedgehog precursor protein (SHhN-SHhC) undergoes a specialized form of autoprocessing called peptide bond cholesterolysis, brought about by the precursor’s enzymatic SHhC domain. A conserved cysteine residue (Cys1) of SHhC initiates the transformation by rearranging the backbone peptide bond at the SHhN-SHhC junction, forming an internal thioester; this SHhN∼SHhC thioester is then displaced by substrate cholesterol, facilitated through general base catalysis involving a conserved aspartate residue (D46). (bottom) Cholestane stereochemistry and numbering system relevant to the present work.

**Figure S1.**
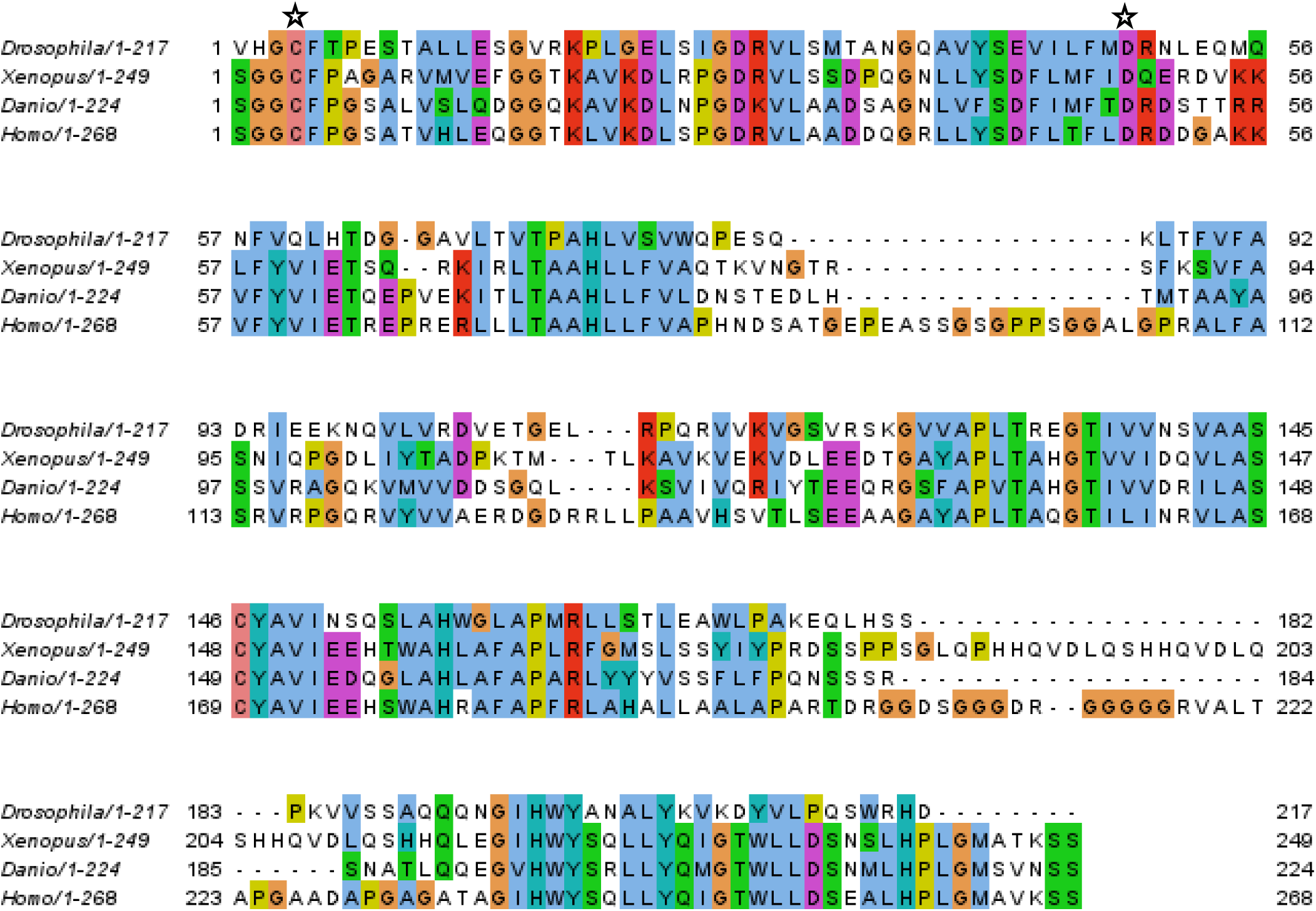
Sequence alignment of *Drosophila*, *Xenopus*, *Danio*, and human SHhC. Uniprot accession numbers: Q02936 (*Drosophila*), Q92008 (*Danio*), Q92000 (*Xenopus*), Q15465 (*Homo*). The identity and similarity values are summarized in **supporting table 1**. The catalytically essential C1 and D46 residues are indicated with a star. Note that the (−3) residue here is numbered as (1).

**Figure S2.**
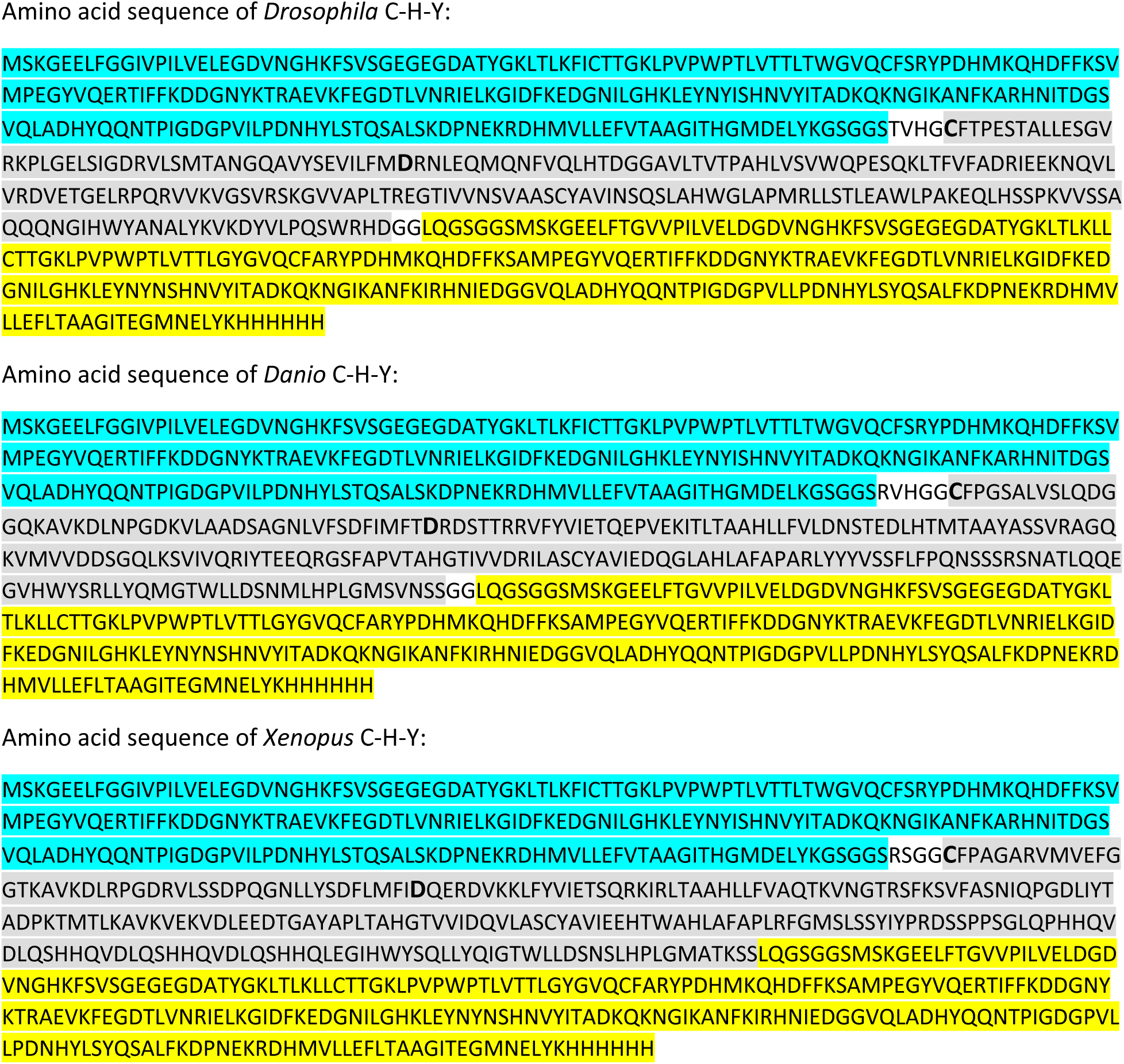
Amino acid sequences of *Drosophila*, *Danio*, and *Xenopus* C-H-Y. Cyan fluorescent protein (cyan); SHhC (gray); yellow fluorescent protein (yellow). The catalytically essential C1 and D46 residues, are labeled in bold. Sequences were verified by whole plasmid sequencing (Azenta).

**Figure S3.**
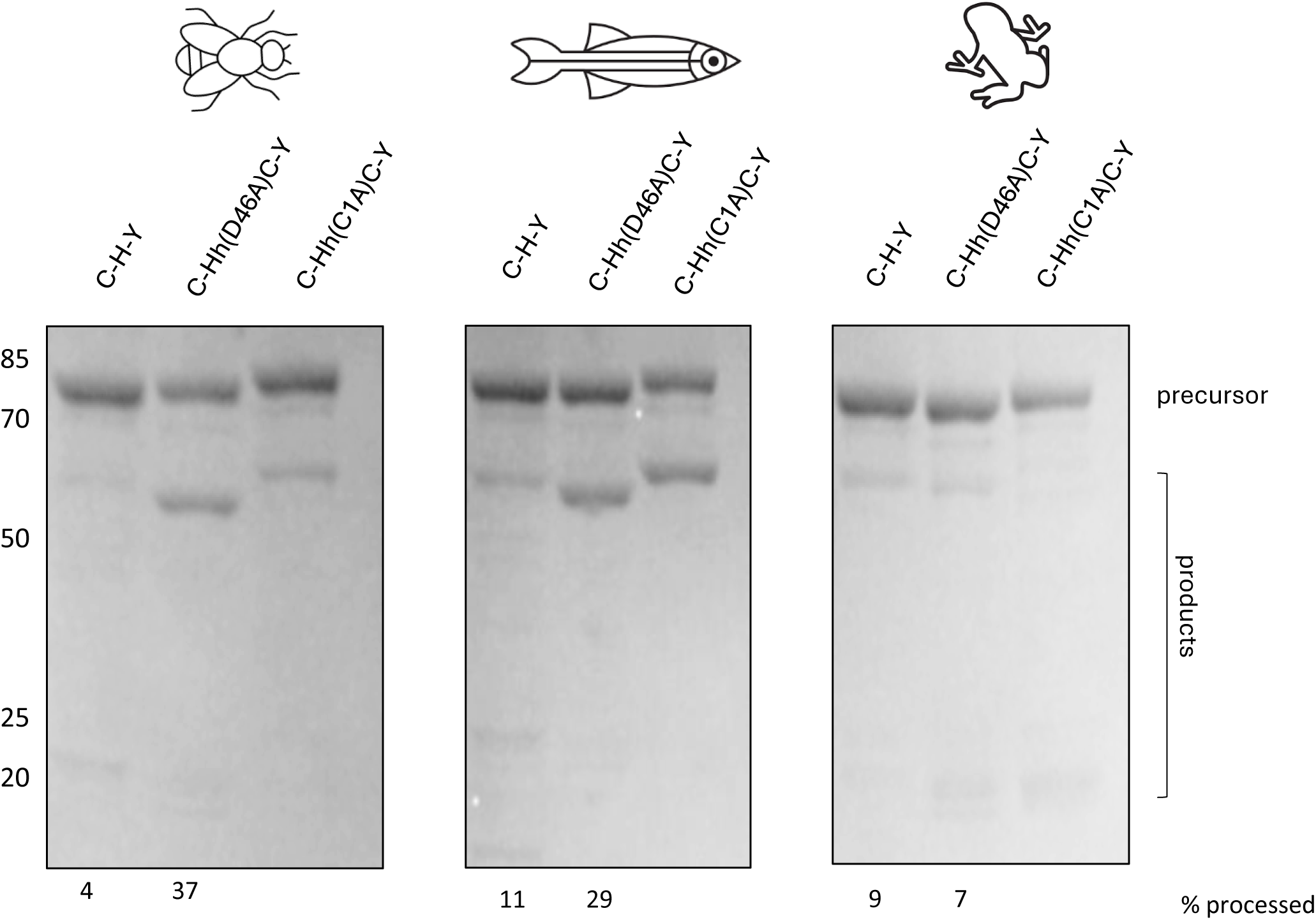
SDS-PAGE of bacterially expressed, Ni-NTA purified Dme, Dre, Xla C-H-Y reporter constructs (WT, D46A, C1A). The extent of precursor autoprocessing that occurred during the expression and purification is indicated below each gel. Protein samples were run at 2 µM precursor concentration, and the extent of processing was determined by ImageJ.

**Figure S4.**
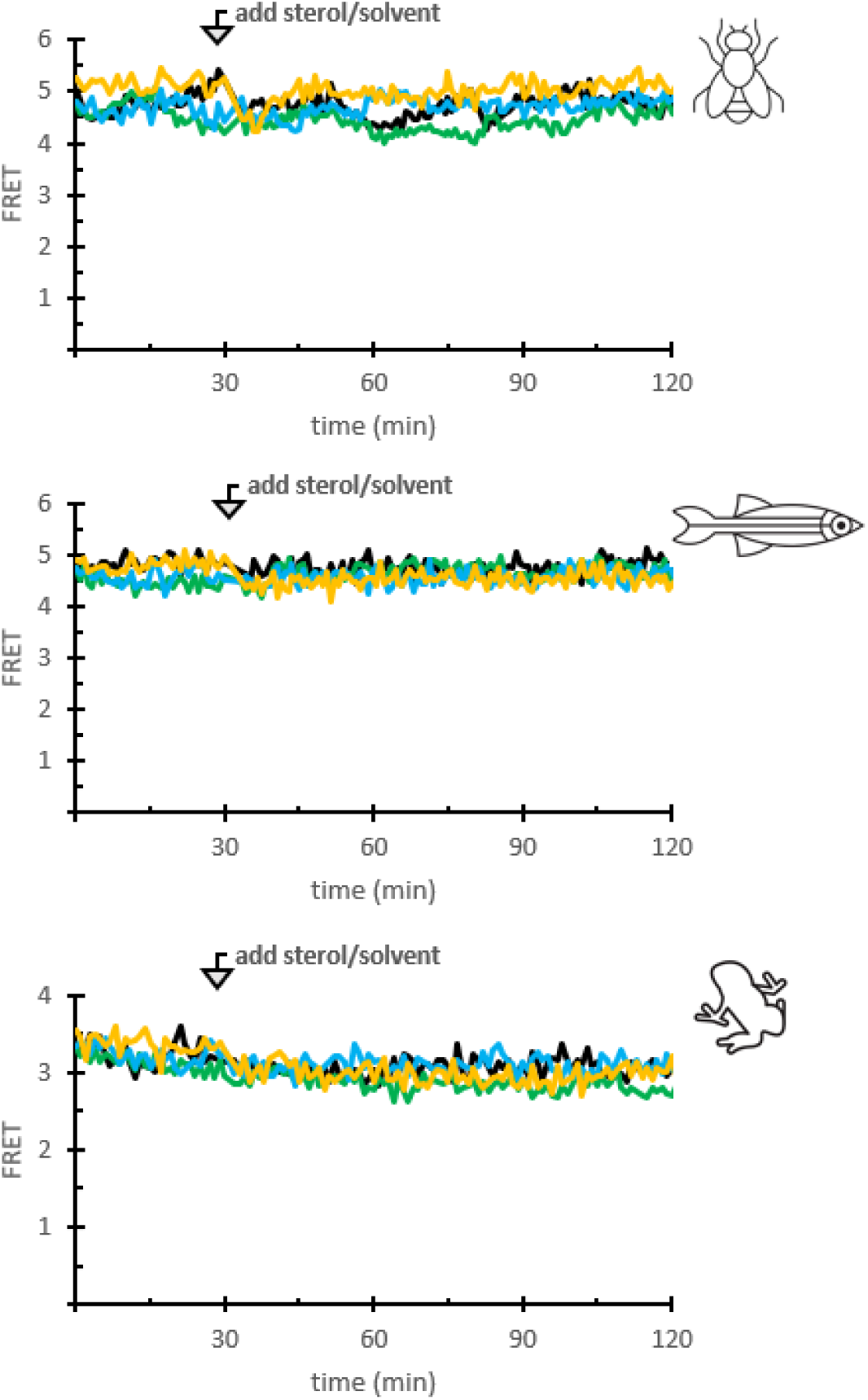
Alanine point mutation at the catalytically essential C1 residue of SHhC eliminates activity. Reactions were run for 2 hours at 30°C in 96-wellplates using 0.2µM C-Hh(C1A)C-Y, Bis-Tris buffer (pH 7.1) containing EDTA (5 mM) and NaCl (0.5 M), and Fos-choline 12 (1.5 mM), with either no sterol (black); 50µM cholesterol (green); 2-ACC (cyan); or 100mM DTT (yellow).

**Figure S5.**
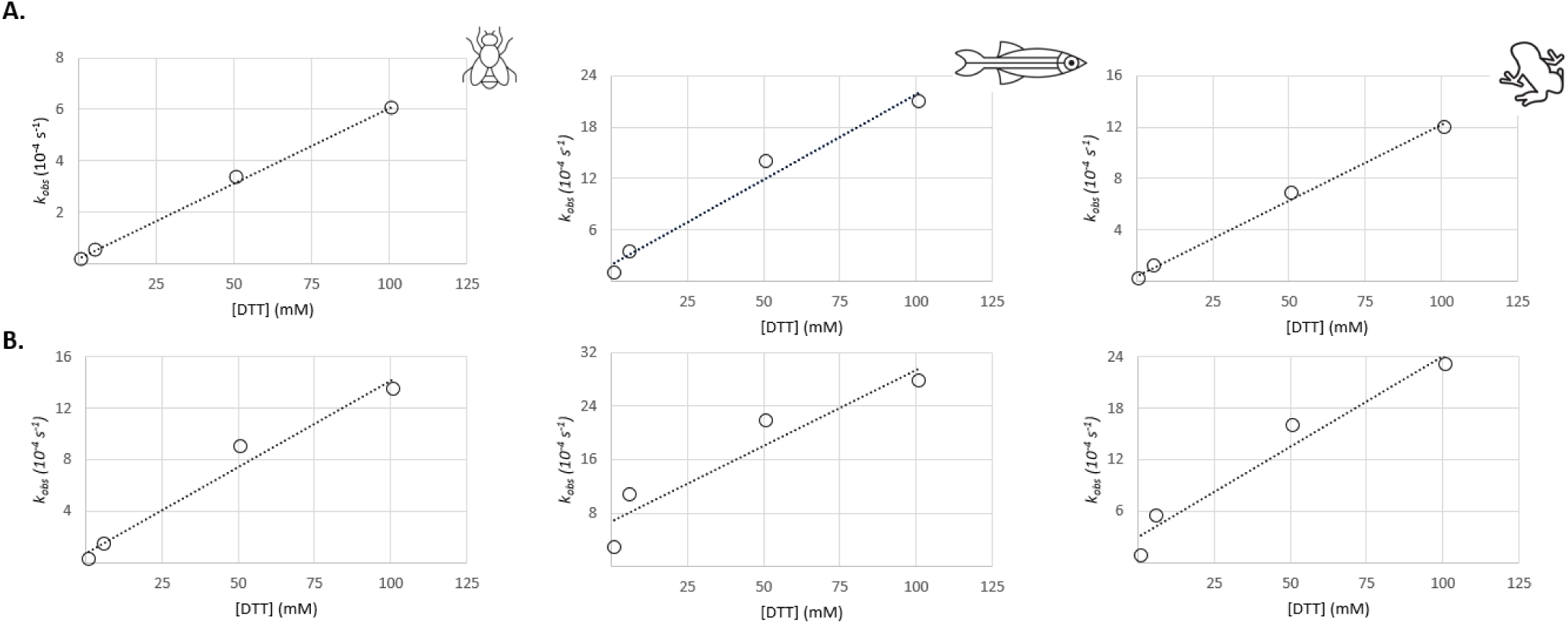
Rates of thioloysis (*k*_obs_) plotted as a function of increasing DTT concentration. Reactions were run for 2 hours at 30°C in 96-well plates using 0.2 µM C-H-Y (*A*) or C-H(D46A)-Y (*B*), Bis-Tris buffer (pH 7.1) also containing EDTA (5mM) and NaCl (0.5M), and Fos-choline 12 (1.5mM). (*A*) DTT concentration vs. *k*_obs_ of DTT-based processing for WT constructs from Dme (left), Dre (middle), or Xla (right). (*B*) DTT concentration vs. *k*_obs_ as an (A), expect for (D46A) mutants.

**Figure S6.**
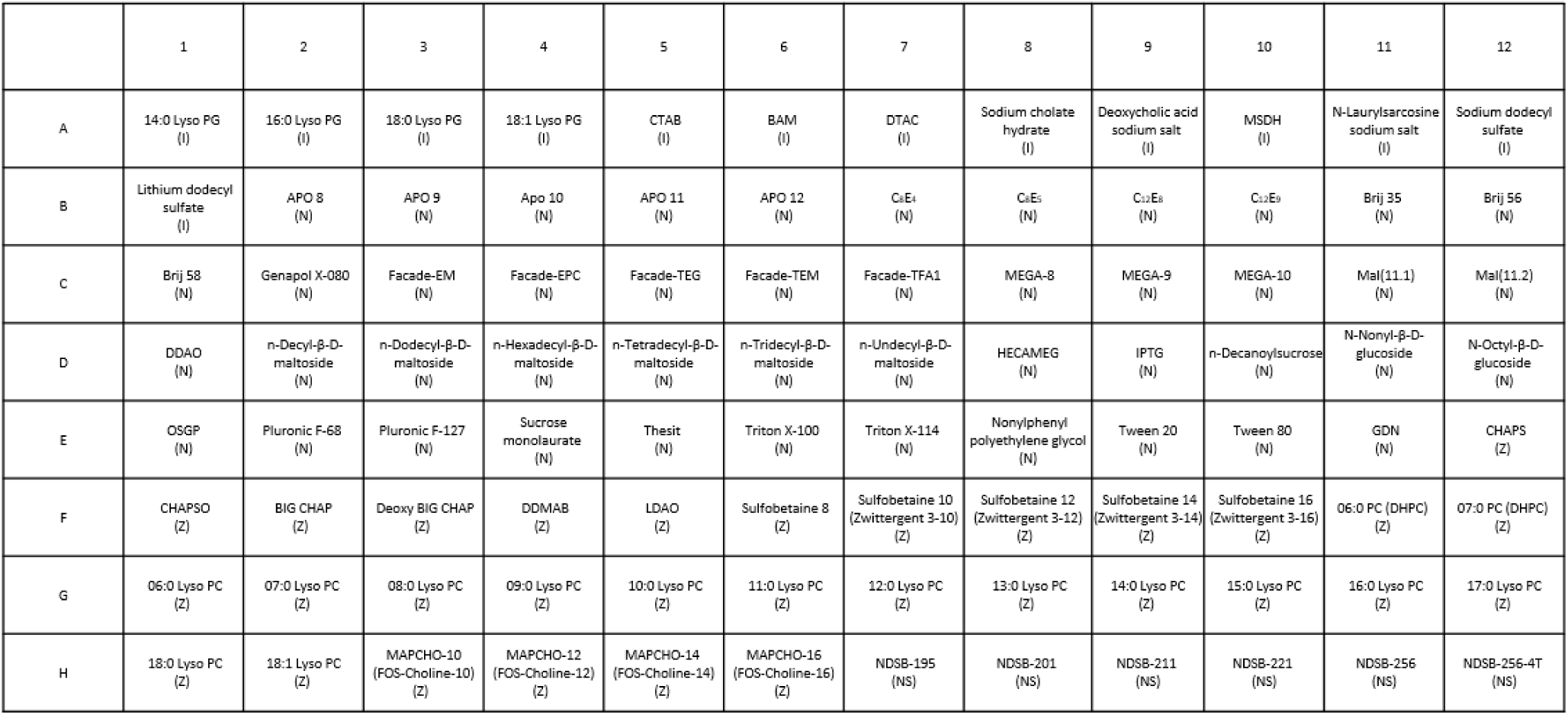
Key for the 96-well detergent screen. This screen from Hampton Research consisted of 13 ionic (I, *wells* **A01-B01**) detergents, 46 non-ionic (N, *wells* **B02-E11**) detergents, 31 zwitterionic detergents (Z, *wells* **E12-H06**), and 6 non-detergent Sulfobetaines (NS, *wells* **H07-H12**). For the ionic detergents, there were 4 cationic (I, *wells* **A05, A06, A07, and A10**), and 9 anionic (I, *wells* **A01-A04, A08, A09, A10-B01**).

**Figure S7.**
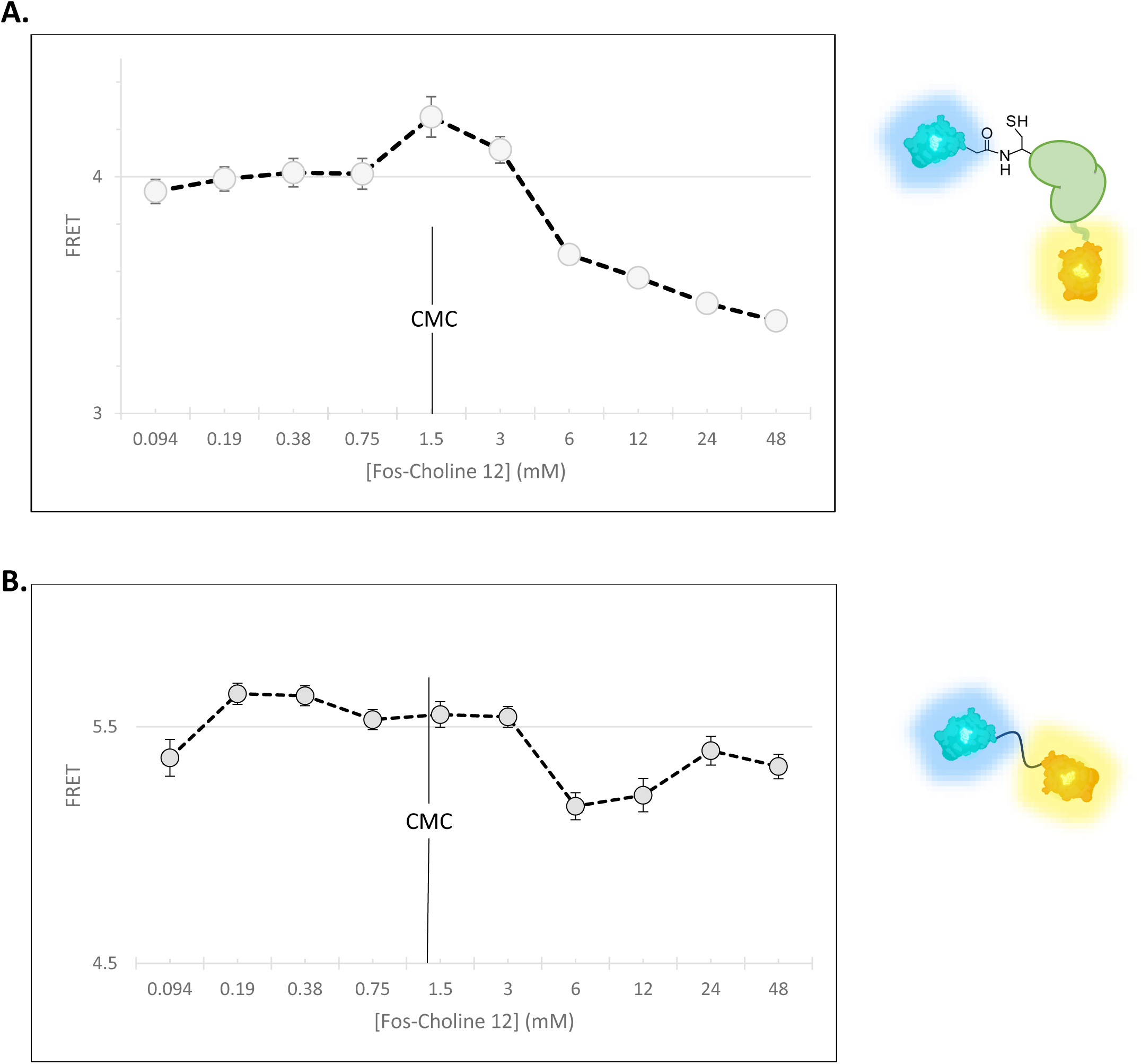
Impact of Fos-Choline 12 concentration on FRET signal from C-H-Y. (*A*) Concentration-response plot of the average FRET for Wild-Type *Drosophila* C-H-Y as a function of increasing Fos-Choline 12. (*B*) Concentration-dependent traces representing the average FRET level recorded from control construct, C-Y, at the same Fos-Choline 12 concentration as in (A).

**Figure S8.**
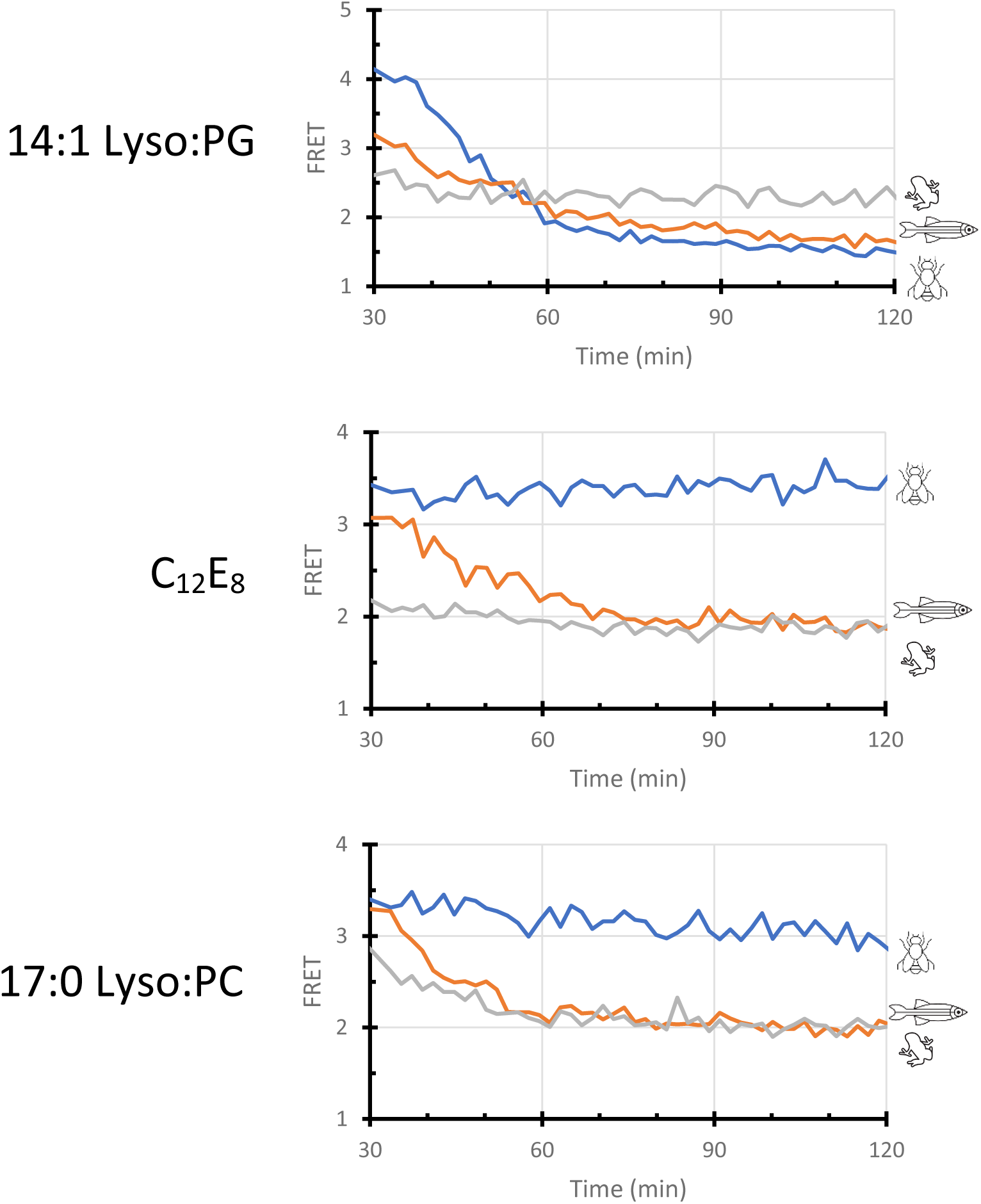
Species specific detergent effects. Kinetic traces for C-H-Y cholesterolysis using either 14:1 Lyso:PG (*well* **A1**, top), C_12_E_8_ (*well* **B9**, middle), or 17:0 Lyso PC (*well* **G12**, bottom) in place of Fos-choline 12. (top) 14:1 Lyso:PG resulted in rapid cholesterolysis for both *Drosophila* (blue) and *Danio* (orange), but no cholesterolysis activity was apparent for *Xenopus* (gray). (middle) C_12_E_8_ resulted in moderate cholesterolysis for *Danio* (orange), but no activity for both *Drosophila* (blue) and *Xenopus* (gray). (bottom) 17:0 Lyso:PC supported rapid cholesterolysis for *Xenopus* (gray), moderate cholesterolysis for *Danio* (orange), and little to no cholesterolysis activity for *Drosophila* (blue). Samples were monitored for 2 hours at 30°C in 96-wellplates using 0.2 µM C-H-Y in Bis-Tris buffer (pH 7.1) with EDTA (5 mM), NaCl (0.5 M), with detergent at the respective CMC.

**Figure S9.**
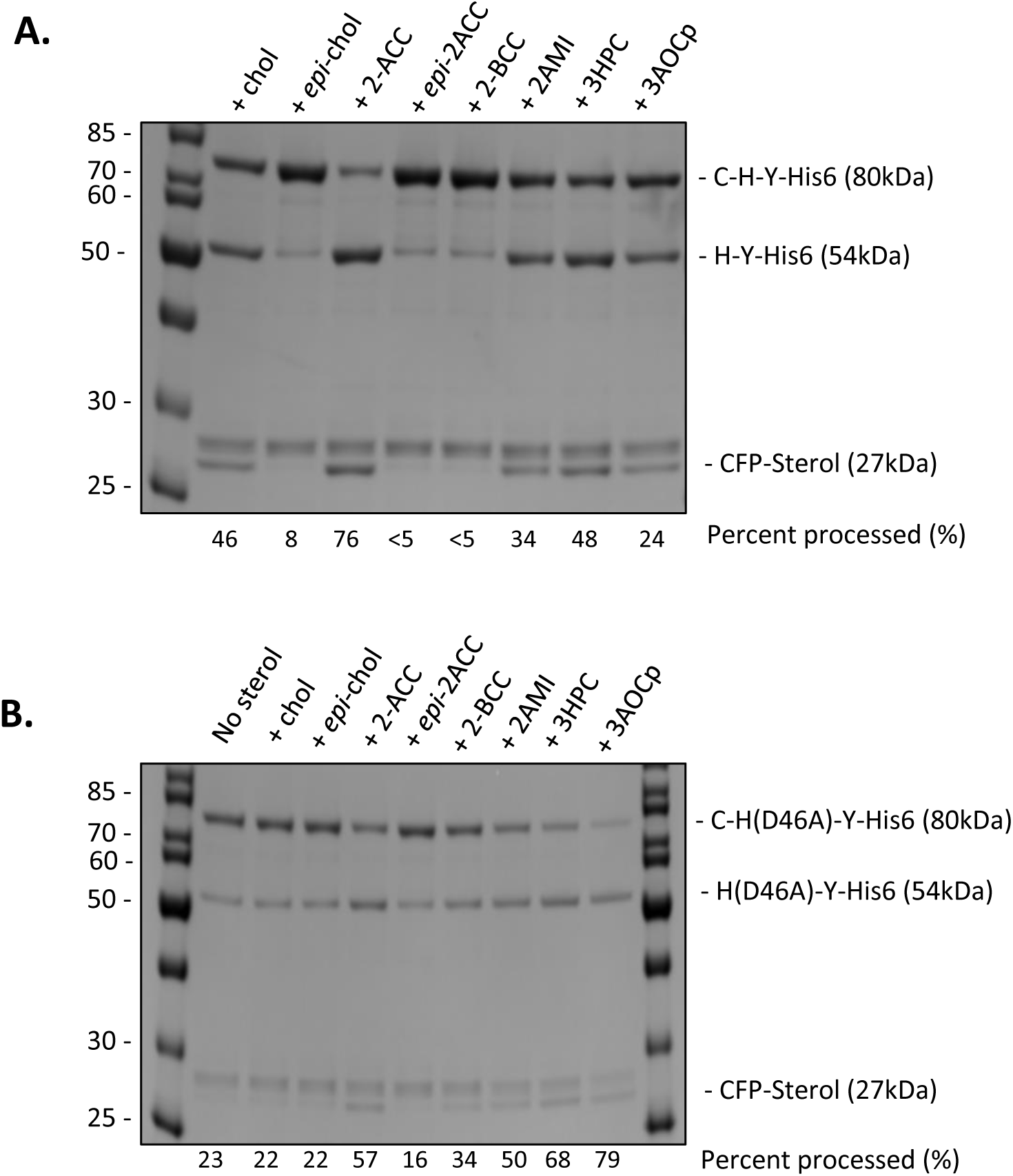
SDS-PAGE Analysis of substrate activity for chemical rescue sterols with *Drosophila* Wild-Type C-H-Y and mutant C-H(D46A)-Y. Reactions were run for 3 hours at 30°C using 1 µM protein, Bis-Tris buffer (pH 7.1) also containing EDTA (5mM) and NaCl (0.5 M), and Fos-choline 12 (1.5 mM). (*A*) SDS-PAGE of *Drosophila* Wild-Type C-H-Y protein with 50 µM of sterol added for each, except 2-BCC which contained 100 µM of sterol. (B) SDS-PAGE of *Drosophila* mutant C-H(D46A)-Y protein with 50 µM of sterol added for each, except 2-BCC which contained 100 µM of sterol. Percent processed values used ImageJ to compare the amount of precursor between the samples (A), or the total percent processing from comparing the product bands to the total band area (B).

**Figure S10.**
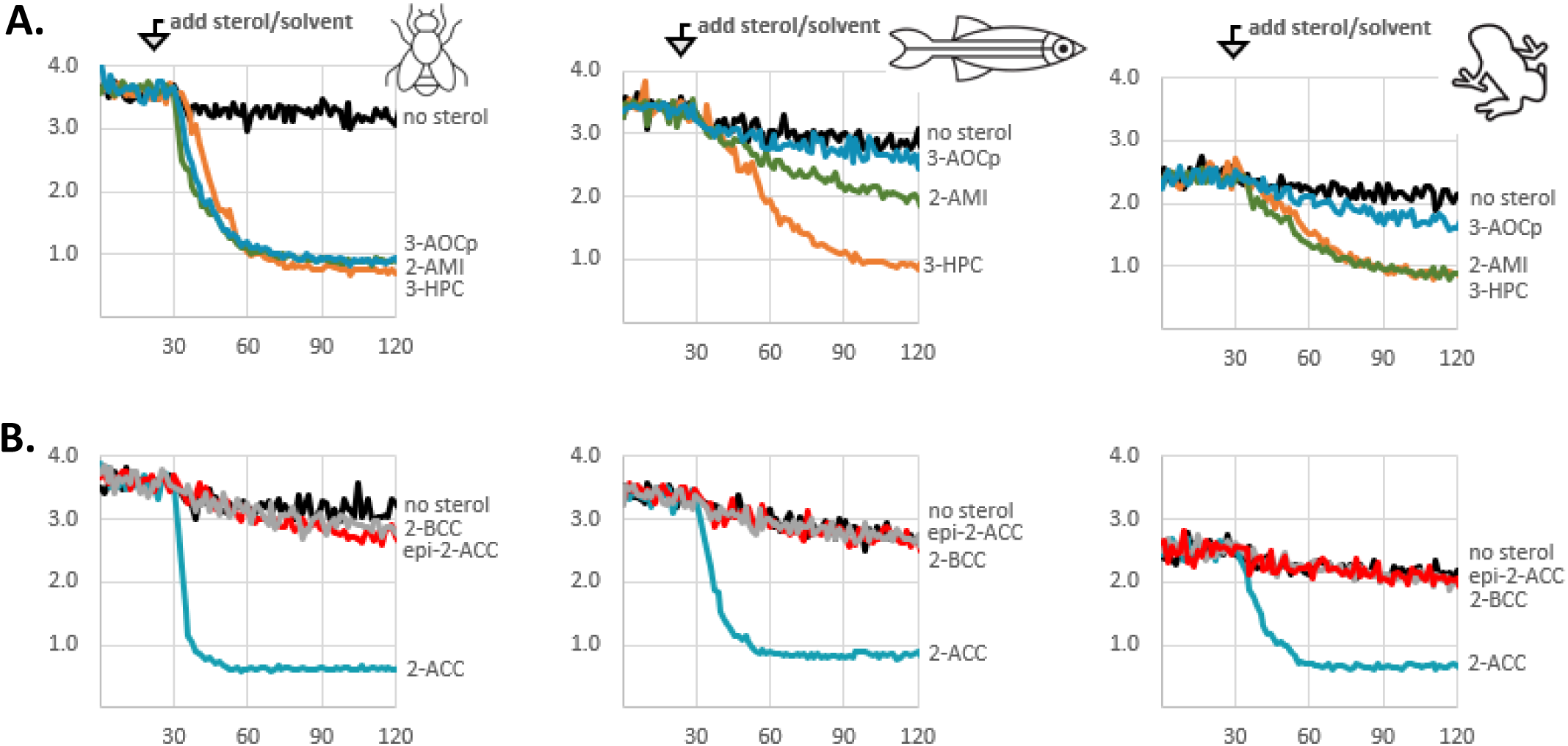
Substrate activity of synthetic sterols using Wild-Type C-H-Y reporter constructs. Samples were monitored for 2 hours at 30°C in 96-wellplates using 0.2 µM C-H-Y in Bis-Tris buffer (pH 7.1) with EDTA (5 mM), NaCl (0.5 M), and Fos-choline 12 (1.5 mM). (*A*) Kinetic traces of *Drosophila* (left), *Danio* (middle), and *Xenopus* (right) C-H-Y with either no sterol (black) or with 50 µM 2-AMI (green), 3-HPC (orange), or 3-AOCp (blue). (*B*) Kinetic traces of *Drosophila* (left), *Danio* (middle), and *Xenopus* (right) C-H-Y with either no sterol (black) or with 50 µM 2-ACC (cyan), 2-BCC (gray), and *epi* 2-ACC (red).

**Figure S11.**
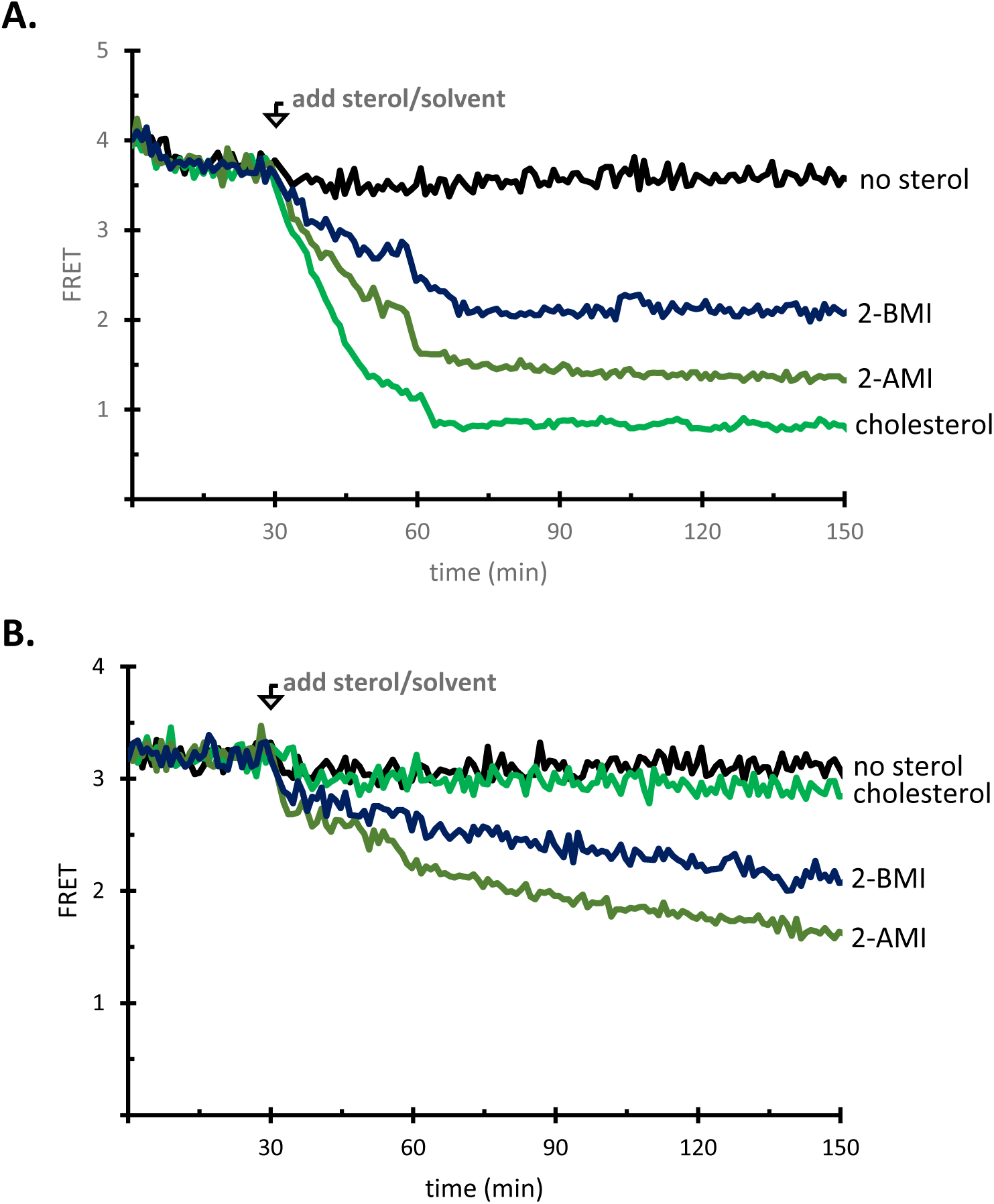
Substrate activity of cholestanol 2-imidazolmethyl epimers with *Drosophila* Wild-Type C-H-Y and mutant C-H(D46A)-Y. Reactions were monitored for 2.5 hours at 30°C in 96-wellplates using 0.2 µM reporter protein in Bis-Tris buffer (pH 7.1) with EDTA (5 mM) and NaCl (0.5 M), and Fos-choline 12 (1.5 mM). (*A*) Kinetic trace of *Drosophila* Wild-Type C-H-Y with either no sterol (black) or with 50 µM cholesterol (light green), 2-AMI (Dark green), or 2-BMI (Blue). (*B*) Kinetic trace of *Drosophila* mutant C-H(D46A)-Y with either no sterol (black) or with 50µM cholesterol (light green), 2-AMI (Dark green), or 2-BMI (Blue).

**Table S1.** Sequence identity and similarity of the cholesterolysis domains from selected model organisms compared to human SHhC^a^.

|  | Dme HhC | Dre SHhC | Xla SHhC |
| --- | --- | --- | --- |
| Identity | 32% | 46% | 49% |
| Similarity | 45% | 60% | 59% |

<sup>a</sup> Identity and similarity were found by Blast2 sequence alignment using SHhC sequences from UniProt, and cited in **supporting figure 1**

**Table S2.** Apparent rates of spontaneous precursor hydrolysis at 30 °C, pH 7.1, for *Drosophila* (Dme), *Danio* (Dre), and *Xenopus* (Xla) Wild-Type and D46A HhC using the FRET reporter C-H-Y.

| Construct | $k_{\max}$<br>(s<sup>-1</sup>) | $t_{1/2}$<br>(hrs) |
| --- | --- | --- |
| Dme HhC (WT) | 0.52*10<sup>-5</sup> | 37 |
| Dre HhC (WT) | 5.0*10<sup>-5</sup> | 3.9 |
| Xla HhC (WT) | 3.2*10<sup>-5</sup> | 6.0 |
| Dme HhC (D46A) | 0.65*10<sup>-5</sup> | 30 |
| Dre HhC (D46A) | 5.4*10<sup>-5</sup> | 3.6 |
| Xla HhC (D46A) | 4.4*10<sup>-5</sup> | 4.4 |

**Table S3.** Thiolysis activity with Dithiothreitol (DTT) for Dme, Dre, and Xla WT and D46A SHhC using FRET reporter.

| Construct | $k_{\max}$<br>($s^{-1}$) | | | |
| --- | --- | --- | --- | --- |
|  | 100mM DTT | 50mM DTT | 5mM DTT | 1mM DTT |
| Dme HhC (WT) | $6.0 \times 10^{-4}$ | $3.4 \times 10^{-4}$ | $0.53 \times 10^{-4}$ | $0.16 \times 10^{-4}$ |
| Dre HhC (WT) | $21 \times 10^{-4}$ | $14 \times 10^{-4}$ | $3.5 \times 10^{-4}$ | $0.93 \times 10^{-4}$ |
| Xla HhC (WT) | $12 \times 10^{-4}$ | $6.9 \times 10^{-4}$ | $1.2 \times 10^{-4}$ | $<0.32 \times 10^{-4}$ |
| Dme HhC (D46A) | $14 \times 10^{-4}$ | $9.0 \times 10^{-4}$ | $1.4 \times 10^{-4}$ | $0.30 \times 10^{-4}$ |
| Dre HhC (D46A) | $28 \times 10^{-4}$ | $22 \times 10^{-4}$ | $11 \times 10^{-4}$ | $2.9 \times 10^{-4}$ |
| Xla HhC (D46A) | $23 \times 10^{-4}$ | $16 \times 10^{-4}$ | $5.5 \times 10^{-4}$ | $0.85 \times 10^{-4}$ |

**Table S4.** Compatible detergents for cholesterolysis with Dme, Dre, and Xla SHhC at 30°C, in Bis-Tris buffer (pH 7.1)

| *Drosophila* |  |  | *Danio* |  |  | *Xenopus* |  |  |
| --- | --- | --- | --- | --- | --- | --- | --- | --- |
| Detergent<sup>a</sup> | Type | $k_{\text{obs}}$<br>(s<sup>-1</sup>) | Detergent<sup>a</sup> | Type | $k_{\text{obs}}$<br>(s<sup>-1</sup>) | Detergent<sup>a</sup> | Type | $k_{\text{obs}}$<br>(s<sup>-1</sup>) |
| 14:0 Lyso PG | I | 0.79*10<sup>-3</sup> | 14:0 Lyso PG | I | 0.56*10<sup>-3</sup> | Brij 35 | N | 0.89*10<sup>-3</sup> |
| 18:0 Lyso PG | I | 0.65*10<sup>-3</sup> | 18:1 Lyso PG | I | 0.65*10<sup>-3</sup> | Brij 56 | N | 0.82*10<sup>-3</sup> |
| Brij 35 | N | 0.69*10<sup>-3</sup> | APO 12 | N | 0.50*10<sup>-3</sup> | Brij 58 | N | 0.89*10<sup>-3</sup> |
| Genapol X-080 | N | 0.80*10<sup>-3</sup> | Brij 56 | N | 0.65*10<sup>-3</sup> | Genapol X-080 | N | 0.85*10<sup>-3</sup> |
| n-Decanoylsucrose | N | 0.89*10<sup>-3</sup> | Brij 58 | N | 0.51*10<sup>-3</sup> | DDAO | N | 0.61*10<sup>-3</sup> |
| Nonylphenyl<br>polyethylene glycol | N | 0.95*10<sup>-3</sup> | Genapol X-080 | N | 0.84*10<sup>-3</sup> | n-Decanoylsucrose | N | 0.68*10<sup>-3</sup> |
| 10:0 Lyso PC | Z | 0.73*10<sup>-3</sup> | Mal(11.2) | N | 0.62*10<sup>-3</sup> | Nonylphenyl<br>polyethylene glycol | N | 1.0*10<sup>-3</sup> |
| 12:0 Lyso PC | Z | 1.2*10<sup>-3</sup> | n-Decyl-β-D-maltoside | N | 0.65*10<sup>-3</sup> | Tween 20 | N | 1.0*10<sup>-3</sup> |
| 13:0 Lyso PC | Z | 0.95*10<sup>-3</sup> | n-Decanoylsucrose | N | 0.63*10<sup>-3</sup> | Tween 80 | N | 1.0*10<sup>-3</sup> |
| 15:0 Lyso PC | Z | 0.93*10<sup>-3</sup> | N-Nonyl-β-D-glucoside | N | 0.74*10<sup>-3</sup> | Zwittergent 3-16 | N | 0.82*10<sup>-3</sup> |
| Fos-Choline 10 | Z | 0.66*10<sup>-3</sup> | Nonylphenyl<br>polyethylene<br>glycol | N | 0.71*10<sup>-3</sup> | 10:0 Lyso PC | Z | 0.73*10<sup>-3</sup> |
| Fos-Choline 12 | Z | 1.9*10<sup>-3</sup> | Tween 20 | N | 0.82*10<sup>-3</sup> | 12:0 Lyso PC | Z | 1.3*10<sup>-3</sup> |
| Fos-Choline 14 | Z | 1.3*10<sup>-3</sup> | Zwittergent 3-16 | N | 1.0*10<sup>-3</sup> | 13:0 Lyso PC | Z | 1.0*10<sup>-3</sup> |
| Fos-Choline 16 | Z | 1.0*10<sup>-3</sup> | 07:0 PC (DHPC) | Z | 1.1*10<sup>-3</sup> | 15:0 Lyso PC | Z | 0.86*10<sup>-3</sup> |

<sup>a</sup> Detergents plate designations can be found in **supporting figure 6**

**Table S4 Continued.**
| *Danio* |  |  | *Xenopus* |  |  |
| --- | --- | --- | --- | --- | --- |
| Detergent<sup>a</sup> | Type | $k_{\text{obs}}$<br>(s<sup>-1</sup>) | Detergent<sup>a</sup> | Type | $k_{\text{obs}}$<br>(s<sup>-1</sup>) |
| 12:0 Lyso PC | Z | $0.98 \cdot 10^{-3}$ | 17:0 Lyso PC | Z | $1.1 \cdot 10^{-3}$ |
| 13:0 Lyso PC | Z | $0.88 \cdot 10^{-3}$ | 18:0 Lyso PC | Z | $1.3 \cdot 10^{-3}$ |
| 15:0 Lyso PC | Z | $0.80 \cdot 10^{-3}$ | 18:1 Lyso PC | Z | $1.1 \cdot 10^{-3}$ |
| 18:0 Lyso PC | Z | $0.84 \cdot 10^{-3}$ | Fos-Choline 10 | Z | $0.99 \cdot 10^{-3}$ |
| 18:1 Lyso PC | Z | $1.2 \cdot 10^{-3}$ | Fos-Choline 12 | Z | $1.6 \cdot 10^{-3}$ |
| Fos-Choline 12 | Z | $1.4 \cdot 10^{-3}$ | Fos-Choline 14 | Z | $0.95 \cdot 10^{-3}$ |
| Fos-Choline 14 | Z | $0.93 \cdot 10^{-3}$ | Fos-Choline 16 | Z | $1.1 \cdot 10^{-3}$ |

<sup>a</sup> Detergents plate designations can be found in **supporting figure 6**

**Table S5.**
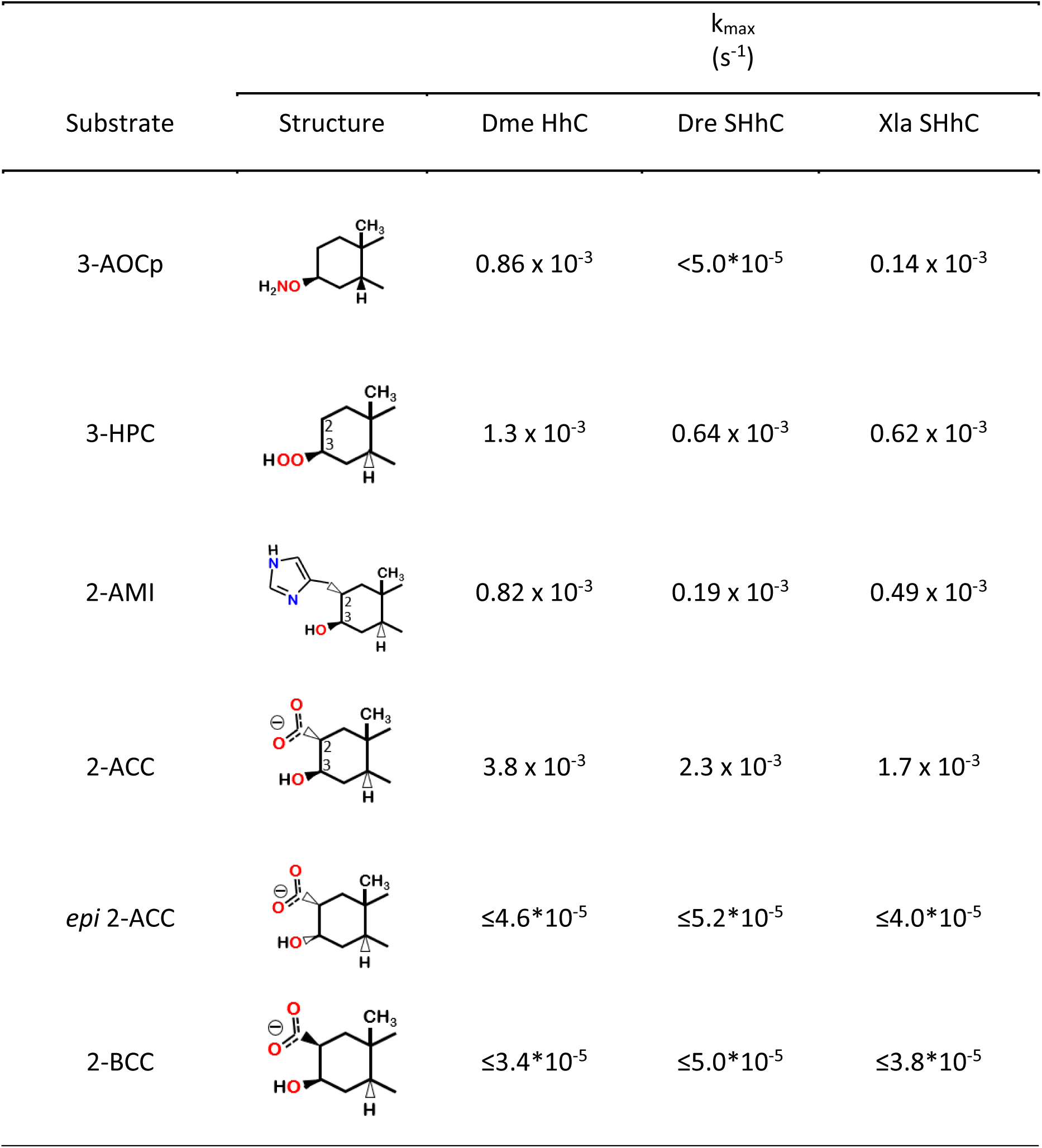
Substrate activity of engineered sterols with *Drosophila*, *Danio*, and *Xenopus* Wild-Type HhC.

| Substrate | Structure | $k_{\max}$<br>($s^{-1}$) | | |
| --- | --- | --- | --- | --- |
|  |  | Dme HhC | Dre SHhC | Xla SHhC |
| 3-AOCp | | $0.86 \times 10^{-3}$ | $<5.0 \times 10^{-5}$ | $0.14 \times 10^{-3}$ |
| 3-HPC | | $1.3 \times 10^{-3}$ | $0.64 \times 10^{-3}$ | $0.62 \times 10^{-3}$ |
| 2-AMI | | $0.82 \times 10^{-3}$ | $0.19 \times 10^{-3}$ | $0.49 \times 10^{-3}$ |
| 2-ACC | | $3.8 \times 10^{-3}$ | $2.3 \times 10^{-3}$ | $1.7 \times 10^{-3}$ |
| *epi* 2-ACC | | $\leq 4.6 \times 10^{-5}$ | $\leq 5.2 \times 10^{-5}$ | $\leq 4.0 \times 10^{-5}$ |
| 2-BCC | | $\leq 3.4 \times 10^{-5}$ | $\leq 5.0 \times 10^{-5}$ | $\leq 3.8 \times 10^{-5}$ |

## Supporting Methods

### Construction of FRET reporter constructs for Xla and Dru SHhC

Gene fragments encoding SHhC with an upstream SHhN peptide were synthesized with 5’ (Xho I) and 3’ (Pst I) sites, codon-optimized for *E. coli* expression, and cloned into a modified pBAD33 FRET expression vector, described previously.^1, 2^ Reporter constructs for wild type, C1A and D46A SHhC were prepared in this same way. Whole plasmid sequencing (Plasmid-EZ, Azenta) was used to confirm construct assembly. The expressed C-H-Y polyprotein carries a C-terminal His-tag for purification.

### *E. coli* expression and Ni-NTA purification of C-H-Y FRET reporter

E. coli strain LMG194 was used mainly for expressing C-H-Y. Following previous work,^3, 4^ a swath of single colonies of the C-H-Y expression strain was transferred from an LB-agar plate containing 50 μg/mL chloramphenicol (CAM) into a 125 mL baffled flask containing 50 mL of LB with CAM (50 μg/mL). The flask was incubated at 37 °C with shaking (225 RPM) until the optical density reading (OD_600_) reached between 0.6-0.8 units. Protein expression was then induced by adding arabinose (0.2 % w/v) and the flask was incubated for additional 18-22 hours at reduced temperature (16 °C). After the induction period, the broth was transferred to 50 mL conical tube and centrifuged at 10,000 RPM for 10 minutes at 4 °C. The broth was discarded and the pellet resuspended in 1 mL of lysis buffer (20mM Na_2_HPO_4_ buffer pH 7, with 400mM NaCl, 100mM KCl, 10mM imidazole, 0.5% Triton X-100, and 10% glycerol). Lysozyme was added to the homogenate (0.2mg / mL, final) and incubated for 10 minutes at room temperature, followed by two freeze-thaw cycles at −80 C. To the viscous mixture, we then added 10 μL of Longlife DNase (G-Biosciences) plus 5 μL of mung bean nuclease (NEB) and incubated at room temperature with gentle mixing for 15 minutes. All steps past this point were done on ice with pre-chilled buffers. Once the total lysate was free flowing, it was combined with an equal volume of 2x binding buffer (40mM Na_2_HPO_4_ pH 7, 1M NaCl, 20mM imidazole, and 20% glycerol). After a final vortex, insoluble cellular debris was removed by centrifugation (14,000 RPM, 15 min). His-tagged C-H-Y was purified from this soluble fraction using a His SpinTrap column (Cytiva). After passing 500 µl of wash buffer 1 (500mM NaCl, 20mM Na_2_HPO_4_, 37.5mM imidazole and 10% glycerol), and 500 µl of wash buffer 2 (500mM NaCl, 20mM Na_2_HPO_4_, 75mM imidazole and 10% glycerol), the reporter protein was eluted with 300 μL of elution buffer (500mM NaCl, 20mM Na_2_HPO_4_, 500 mM imidazole and 10% glycerol). To protect cysteine residues of SHhC, 3 μL of TCEP (500mM) was added to the elution and stored at −80 C. Concentration and purity of C-H-Y in the elution was determined by SDS-PAGE using BioRad Gel Doc EZ system.

### FRET assays for SHhC cholesterolysis

Activity measurements were conducted in 100 ul volume with C-H-Y between 100-200 nM, along with Bis-Tris propane buffer (50 mM, pH 7.2), Ethylenediaminetetraacetic acid (EDTA) (0.5 mM) and NaCl (100 mM). Unless stated otherwise, we used Fos-choline 12 (1.5 mM, final) to solubilize cholesterol and TCEP (5 mM, final) as a reducing agent. Assays were conducted at 30 °C in 96 well NBS™ Microplates (Corning) using a BioTek Synergy H1 plate reader with Gen5 software. Sample FRET readings were recorded every 1-2 minutes as the 540nm/460nm emission ratio after excitation at 460 nm. Reactions were initiated by addition of cholesterol from an ethanol stock.

### Determination of k_max_ and K_M_ values from FRET assay data

For each C-H-Y/sterol, the maximum rate of sterolysis (k_max_) was estimated from the experimental FRET loss data at the highest sterol concentration by curve fitting to a first order exponential decay.

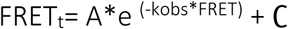

K_M_ values for substrate sterols in the C-H-Y system was estimated by plotting the initial rate of FRET loss (slope of experimental data) as a function of increasing initial substrate concentration. Curve fitting to a Michaelis-Menten equation was carried out using GraphPad prism and Excel.

### Chemical Rescue

Kinetic experiments were carried using the FRET approach (as above) except that chemically modified cholestanol derivatives replaced cholesterol as potential substrates. FRET assays included six replicates for each C-H-Y construct (WT and D46A) and each cholestanol derivative. Selected compounds, 2-ACC and 2-BCC, were tested further in concentration-response experiments (as above) to determine K_M_ value.

### Detergent screening for compatibility with SHhC cholesterolysis

Detergent screening for Dme, Xla, and Dre cholesterolysis used the 96-member library from Hampton Research (cat: HR2407). In preparation for the screen, the detergent library was thawed at room temperature, then warmed to 37°C until all compounds had fully dissolved. Once complete, 10 μL of each detergent was transferred to the 96-well assay plate, followed by the addition of 88 uL of master mix containing C-H-Y in the buffer components used in the cholesterolysis assays (above), with the exception of fos-choline 12. Prior to initiating cholesterolysis, FRET readings from each sample were recorded for 30 minutes for the purpose of isolating potential detergent effects on C-H-Y FRET signal. After that pre-incubation, cholesterolysis was initiated by adding 2 μL of 2.5 mM cholesterol stock to each well and FRET was monitored for another 1.5 hours. Reaction progress curves for FRET loss were analyzed for selected wells to obtain the apparent k_max_ values.

### Thiolysis of SHhC precursor using the FRET reporter

C-H-Y thiolysis assays were carried out in a manner similar to the cholesterolysis experiments except that here the reactions were initiated with dithiothreitol (DTT) added from aqueous stock in Bis-Tris buffer. First order exponential decay curves were fit to experimental FRET-loss data to determine k_DTT_ values.

### General Procedures

NMR spectra were acquired using a Bruker Avance III 600 MHz or 800 MHz spectrometer at 25 ^°^C. Calibration was by the residual solvent signal (CDCl_3_: ^1^H = 7.26 ppm. ^13^C = 77.0 ppm). Preparative TLC was performed on glass-backed plates (10 cm in length) coated with a 0.25 mm layer of silica gel 60 F254.

### Synthesis of 2-carboxycholestanol isomers

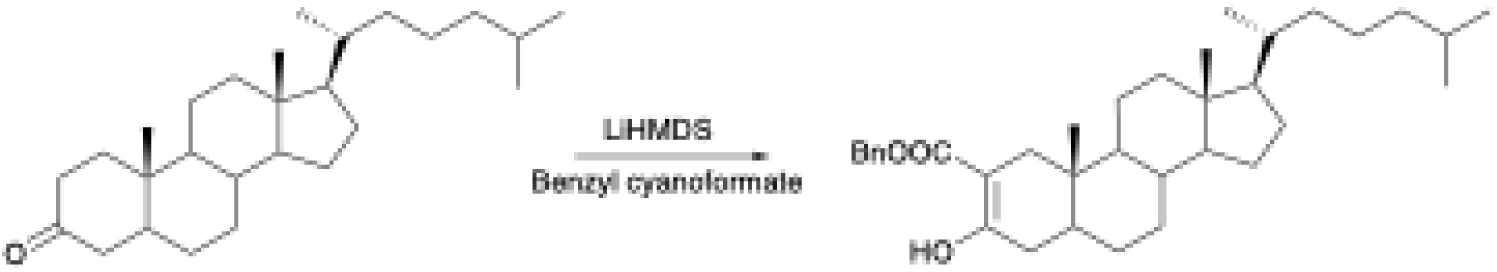

A solution of 5α-cholestan-3-one in anhydrous THF (75 mg/mL) under N_2_ at −78 ^°^C was treated with LiHMDS (1.25 M in THF, 1.3 eq), and the solution was stirred for 20 min., followed by the addition of benzyl cyanoformate (1.25 eq.).^5^ The reaction was allowed to warm to rt over two hours, then poured into 30 mL of 5% conc. HCl and extracted with 3 x 15 mL of hexanes/EtOAc 4:1. The combined organic layers were washed with 10% conc. HCl and brine, then dried over Na_2_SO_4_ and concentrated *in vacuo* to yield the product. Minor *O*-acyl and 4-acyl byproducts visible by NMR could not be separated, and the crude product (80% pure by NMR) was used in the next step without further purification.

2-(Benzyloxycarbonyl)-5α-cholestan-3-one (enol form): ^1^H NMR (600 MHz, CDCl_3_): 12.09 (1H, s), 7.40-7.30 (5H, m), 5.203 (2H, dd, *J =* 12.6, 38.1 Hz), 2.353 (1H, d, *J =* 15.8 Hz), 0.900 (3H, d, *J =* 6.5Hz), 0.864 (3H, d, *J =* 6.6 Hz), 0.860 (3H, d, *J =* 6.6 Hz), 0.747 (3H, s), 0.658 (3H, s).

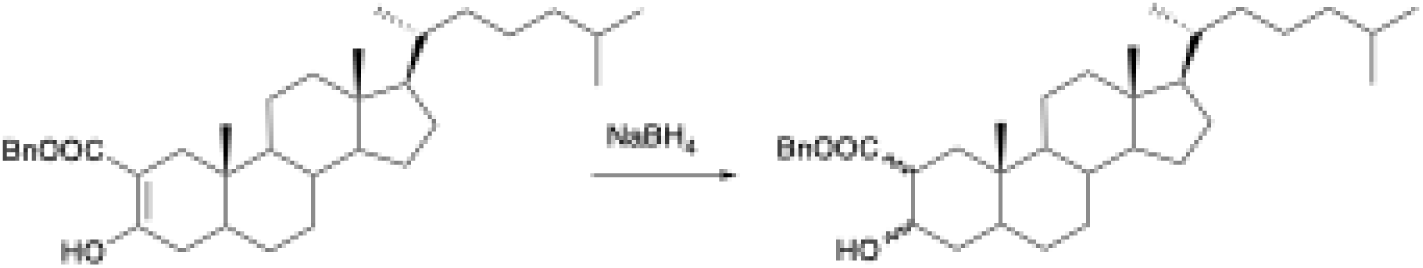

To a solution of the β-keto ester in 2:1 DCM/MeOH (9 mg/mL) was added NaBH_4_ (4 eq) in one portion. The reaction was monitored by TLC, and when no starting material was visible, the reaction was quenched with 5% conc. HCl. The contents were extracted thoroughly with hexanes/EtOAc 4:1, and the combined organic layers were washed with brine, dried over Na_2_SO_4_, and concentrated *in vacuo* to yield the crude product which was purified by preparative TLC (hexanes/EtOAc 4:1).

(2α,3α)-2-Benzyloxycarbonylcholestan-3-ol (24%). ^1^H NMR (600 MHz, CDCl_3_): 7.40-7.31 (5H, m), 5.145 (2H, dd, *J =* 12.3, 29.9 Hz), 4.236 (1H, m), 3.088 (1H, br s), 2.615 (1H, ddd, *J =* 2.3, 3.5, 13.4 Hz), 0.893 (3H, d, *J =* 6.5 Hz), 0.863 (3H, d, *J =* 6.6 Hz), 0.859 (3H, d, *J =* 6.6 Hz), 0.799 (3H, s), 0.643 (3H, s).

(2β,3α)-2-Benzyloxycarbonylcholestan-3-ol (10%). ^1^H NMR (600 MHz, CDCl_3_): 7.40-7.31 (5H, m), 5.131 (2H, dd, *J =* 12.3, 19.3 Hz), 4.063 (1H, m), 3.490 (1H, br s), 2.512 (1H, dd, *J =* 1.7, 12.2 Hz), 0.893 (3H, d, *J =* 6.5 Hz), 0.863 (3H, d, *J =* 6.6 Hz), 0.859 (3H, d, *J =* 6.6 Hz), 0.811 (3H, s), 0.636 (3H, s).

(2α,3β)-2-Benzyloxycarbonylcholestan-3-ol (36%). ^1^H NMR (600 MHz, CDCl_3_): 7.40-7.31 (5H, m), 5.156 (2H, dd, *J =* 12.4, 18.4 Hz), 3.836 (1H, td, *J =* 4.9, 10.7 Hz), 2.614 (1H, br s), 2.529 (1H, ddd, *J =* 3.8, 10.3, 13.1 Hz), 0.892 (3H, d, *J =* 6.5 Hz), 0.862 (3H, d, *J =* 6.6 Hz), 0.858 (3H, d, *J =* 6.6 Hz), 0.839 (3H, s), 0.641 (3H, s).

(2β,3β)-2-Benzyloxycarbonylcholestan-3-ol (30%). ^1^H NMR (600 MHz, CDCl_3_): 7.41-7.33 (5H, m), 5.193 (1H, d, *J =* 12.3 Hz), 5.061 (1H, d, *J =* 12.3 Hz), 3.693 (1H, d, *J =* 11.8 Hz), 3.606 (1H, tt, *J =* 4.8, 11.8 Hz), 2.933 (1H, t, *J =* 5.9 Hz), 2.400 (1H, dd, *J =* 2.3, 14.0 Hz), 1.951 (1H, dt, *J =* 3.3, 12.6 Hz), 0.891 (3H, d, *J =* 6.5 Hz), 0.861 (3H, d, *J =* 6.6 Hz), 0.857 (3H, d, *J =* 6.6 Hz), 0.646 (3H, s), 0.622 (3H, s).

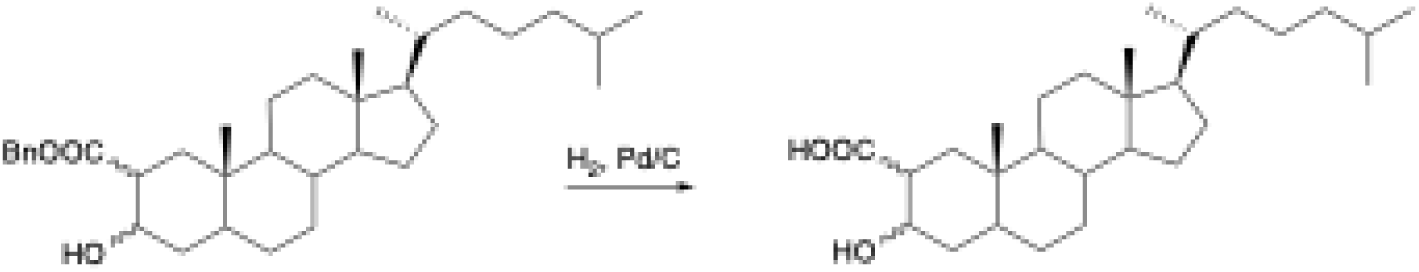

To a solution of each hydroxy ester in EtOAc (5 mg/mL) was added 10% Pd/C (2 mg/mL) and the mixture was stirred under H_2_ (1 atm) for two hrs. CHCl_3_ was added to dissolve the solid products that precipitated during the course of the reaction, followed by filtration through celite to remove the catalyst. Concentration *in vacuo* afforded the deprotected hydroxy acids in quantitative yield.

(2α,3α)-2-Carboxycholestan-3-ol (*epi* 2-ACC). ^1^H NMR (600 MHz, CDCl_3_): 4.272 (1H, m), 2.632 (1H, d, *J =* 13.0 Hz), 0.898 (3H, d, *J =* 6.5 Hz), 0.863 (3H, d, *J =* 6.6 Hz), 0.859 (3H, d, *J =* 6.6 Hz), 0.823 (3H, s), 0.651 (3H, s).

(2β,3α)-2-Carboxycholestan-3-ol (*epi* 2-BCC). ^1^H NMR (600 MHz, CDCl_3_): 4.165 (1H, m), 2.519 (1H, d, *J =* 12.1 Hz), 0.897 (3H, d, *J =* 6.5 Hz), 0.864 (3H, d, *J =* 6.6 Hz), 0.859 (3H, d, *J =* 6.6 Hz), 0.839 (3H, s), 0.649 (3H, s).

(2α,3β)-2-Carboxycholestan-3-ol (2-ACC). ^1^H NMR (600 MHz, CDCl_3_): ^1^H NMR (800 MHz, CDCl_3_): 3.818 (1H, td, *J =* 4.7, 10.6 Hz), 2.513 (1H, td, *J =* 3.0, 11.9 Hz), 2.048 (1H, dd, *J =* 3.2, 13.8 Hz), 1.965 (1H, dt, *J =* 3.0, 12.7 Hz), 1.809 (1H, m), 1.658 (2H, m), 1.58-1.47 (3H, m), 0.896 (3H, d, *J =* 6.4 Hz), 0.864 (3H, d, *J =* 6.6 Hz), 0.860 (3H, d, *J =* 6.6 Hz), 0.855 (3H, s), 0.673 (1H, td, *J =* 3.6, 11.4 Hz), 0.650 (3H, s). ^13^C NMR (201 MHz): 179.57, 71.14, 56.40, 56.23, 54.03, 46.79, 44.50, 42.55, 40.05, 39.87, 39.51, 36.16, 36.07, 35.78, 35.44, 31.84, 28.26, 28.21, 28.01, 24.18, 23.82, 22.81, 22.55, 21.30, 18.67, 12.52, 12.07.

(2β,3β)-2-Carboxycholestan-3-ol (2-BCC). ^1^H NMR (800 MHz, CDCl_3_): 3.715 (1H, dt, *J =* 5.0, 11.9 Hz), 2.937 (1H, t, *J =* 5.7 Hz), 2.477 (1H, dd, *J =* 1.9, 14.1 Hz), 1.972 (1H, dt, *J =* 3.2, 12.8 Hz), 0.899 (3H, d, *J =* 6.6 Hz) 0.866 (3H, d, *J =* 6.6 Hz), 0.862 (3H, d, *J =* 6.6 Hz), 0.802 (3H, s), 0.642 (3H, s), 0.615 (1H, m). ^13^C NMR (201 MHz): 177.89, 71.34, 56.39, 56.32, 54.64, 46.14, 43.22, 42.63, 40.01, 39.94, 39.52, 36.24, 36.17, 35.77, 35.20, 31.85, 28.20, 28.19, 28.01, 24.17, 23.84, 22.79, 22.55, 21.46, 18.68, 12.73, 12.10.

### Synthesis of 2-imidazolemethylcholestanol isomers

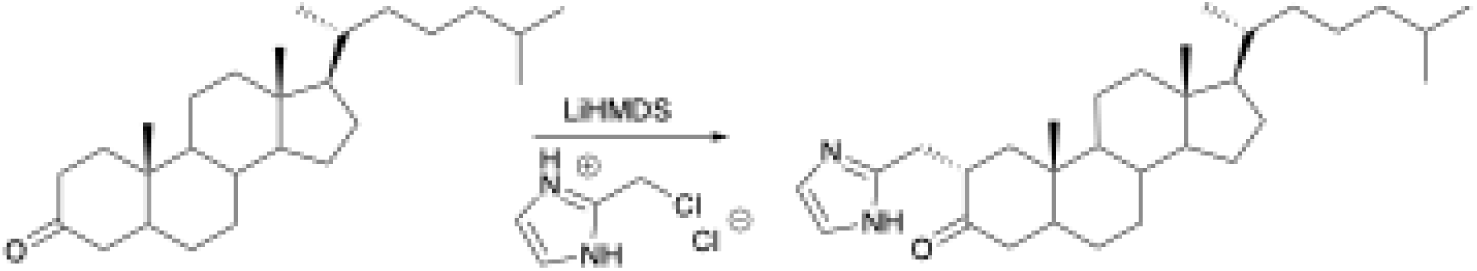

A solution of cholestan-3-one (251 mg, 0.65 mmol) in 5 mL of anhydrous THF under N_2_ was cooled to −78 ^°^C. To this was added 1.3 mL of 1 M LiHMDS/hexane (2.0 eq.) and the solution was stirred for 20 minutes at −78 ^°^C, at which time 2-chloromethyl-1H-imidazole hydrochloride (100 mg, 1.0 eq) was added, followed by in 2 mL of anhydrous THF. The reaction was allowed to slowly warm to rt with stirring for 2.25 hours, then poured into saturated NaHCO_3_ and extracted three times with EtOAc. The combined organic layers were dried over Na_2_SO_4_ and concentrated *in vacuo*. The resulting crude product was largely starting material (*ca.* 60%), but contained *ca.* 30% 2α-product, accompanied by *ca.* 5% each of the 4α- and 2,2-dialkylated products. A sample was subjected to preparative TLC (2% TEA in EtOAc) to isolate the product.

2α-(1H-Imidazol-2-ylmethyl)-cholestan-3-one: ^1^H NMR (600 MHz, CDCl_3_): 6.904 (2H, s), 2.959 (1H, dd, *J =* 9.5, 14.8 Hz), 2.93-2.77 (2H, m), 2.355 (1H, t, *J =* 14.2 Hz), 2.148 (1H, dd, *J =* 5.7, 13.1 Hz), 2.102 (1H, dd, *J =* 3.6, 14.0 Hz), 1.980 (1H, dt, *J =* 3.4, 12.7 Hz), 1.054 (3H, s), 0.898 (3H, d, *J =* 6.6 Hz), 0.865 (3H, d, *J =* 6.6 Hz), 0.860 (3H, d, *J =* 6.6 Hz), 0.664 (3H, s).

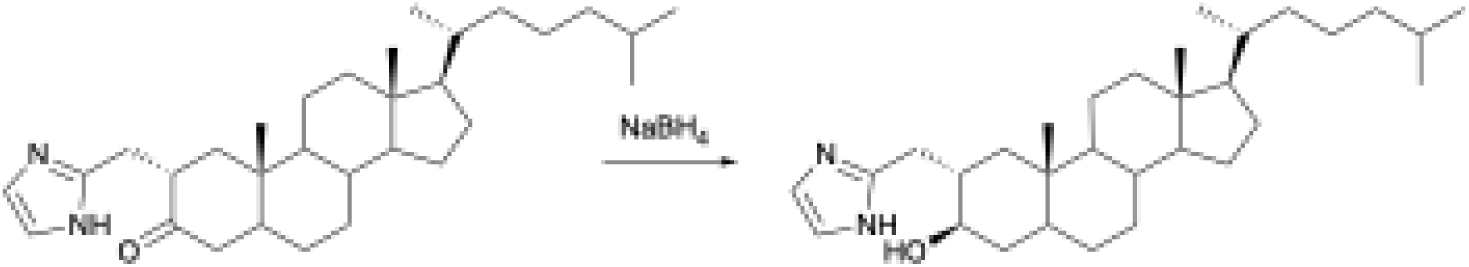

To a solution of imidazole ketone (18.1 mg, 0.039 mmol) in 1.5 ml 4:1 MeOH/DCM, was added CeCl_3_-7H_2_O (24 mg, 1.7 eq.) and the solution was mixed well until the solids had dissolved. An excess of solid NaBH_4_ was then added (10 mg, 6.7 eq) and the solution was stirred for 1.5 hr. The solvent was evaporated to approx. half the volume with a stream of N_2_ and the mixture was extracted with satd. NaHCO_3_ and EtOAc. The combined organic extracts were dried with Na_2_SO_4_ and the solvent evaporated with a stream of N_2_. Purification was accomplished by preparative TLC (1% TEA in EtOAc/MeOH 19:1).

2α-(1H-Imidazol-2-ylmethyl)-cholestan-3β-ol (2-AMI). ^1^H NMR (600 MHz, CDCl_3_): 6.968 (2H, s), 3.358 (1H, td, *J =* 4.6, 10.6 Hz), 2.91-2.82 (2H, m), 0.897 (3H, d, *J =* 6.5 Hz), 0.865 (3H, d, *J =* 6.6 Hz), 0.860 (3H, d, *J =* 6.6 Hz), 0.803 (3H, s), 0.636 (3H, s). ^13^C NMR (201 MHz, CDCl_3_): 147.94, 119.67, 75.33, 56.42, 56.28, 54.18, 44.91, 44.37, 42.56, 39.92, 39.76, 39.51, 38.02, 36.35, 36.16, 35.78, 35.27, 32.10, 31.92, 29.69, 28.22, 28.00, 24.18, 23.84, 22.81, 22.55, 21.25, 18.67, 12.94, 12.07.

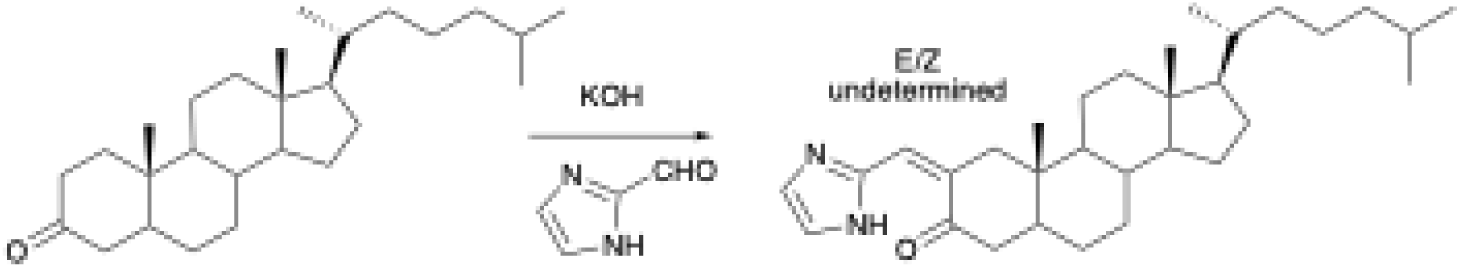

In an alternative synthesis that also provided the 2β-isomer (2-BMI), cholestane-3-one (75.2 mg, 0.19 mmol) in 2.5 ml EtOH was treated with 21 mg powdered KOH (2 eq.) and 22 mg imidazole-2-carbaldehyde (1,2 eq.) at reflux for 5 hr. The mixture was poured into H_2_O containing NaCl and extracted with hexanes/EtOAc 1:2. The organic layers were dried over Na_2_SO_4_ and concentrated *in vacuo*. The product was purified by washing with hexanes.

2-(1H-Imidazol-2-ylmethylene)-cholestan-3β-ol. ^1^H NMR (600 MHz, CDCl_3_): 7.388 (2H, s), 7.242 (1H, s), 3.938 (1H, d, *J =* 17.4 Hz), 2.451 (1H, dd, *J =* 5.3, 19.0 Hz), 2.29-2.20 (2H, m), 2.06-2.02 (1H, m), 0.919 (3H, d, *J =* 6.5 Hz), 0.869 (3H, d, *J =* 6.6 Hz), 0.866 (3H, d, *J =* 6.6 Hz), 0.827 (3H, s), 0.669 (3H, s).

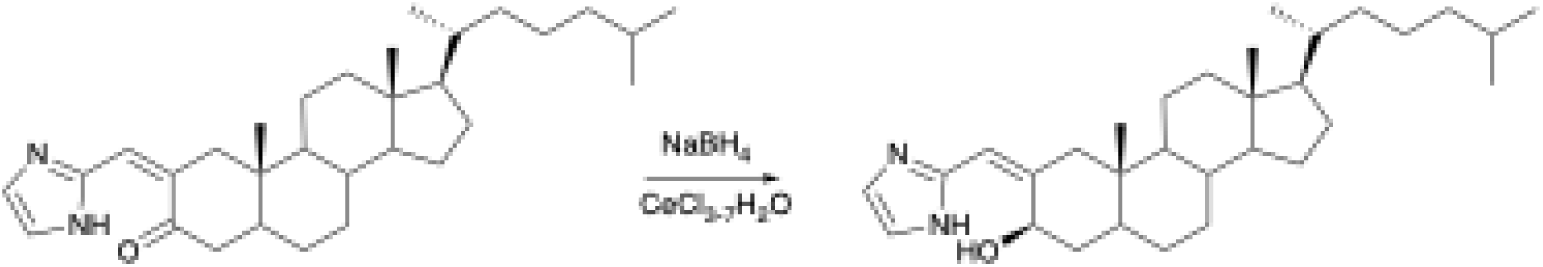

The imidazolylmethylene ketone (4.6 mg, 10 mmol) was dissolved in 1 ml 3:1 MeOH/DCM, an excess of CeCl_3_-7H_2_O (11.1 mg, 3 eq.) was added and stirred for 5 min., followed by addition of NaBH_4_ (excess). After 20 min., the solvent was evaporated to approx. half the volume with a stream of N_2_ and a few drops of 10% HCl were added to destroy the excess NaBH_4_. The mixture was extracted with half-saturated NaHCO_3_ and EtOAc, dried with Na_2_SO_4_, and the solvent was evaporated with a stream of N_2_.

2-(1H-Imidazol-2-ylmethylene)-cholestan-3β-ol. ^1^H NMR (600 MHz, CDCl_3_): 7.069 (2H, s), 6.465 (1H, s), 4.18-4.13 (1H, m), 4.18-4.13 (1H, m), 3.500 (1H, d, *J =* 13.1 Hz), 0.893 (3H, d, *J =* 6.4 Hz), 0.861 (3H, d, *J =* 6.6 Hz), 0.857 (3H, d, *J =* 6.6 Hz), 0.609 (3H, s), 0.605 (3H, s).

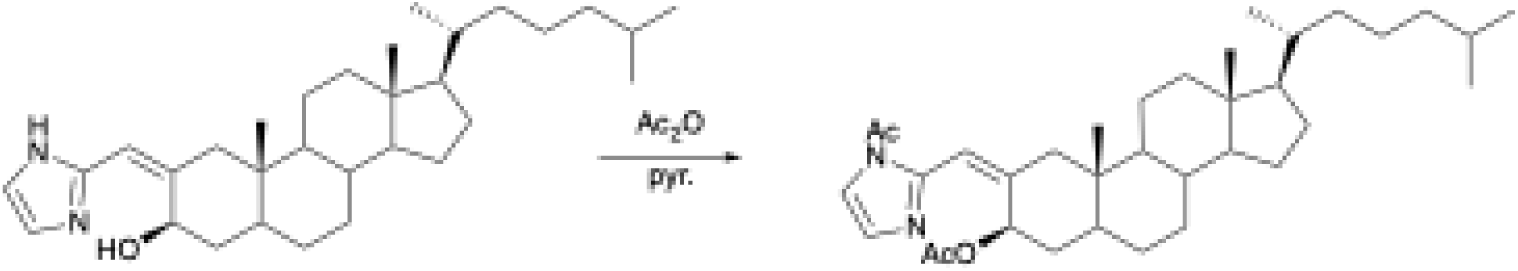

The imidazolylmethylene sterol was acetylated using 3 drops of Ac_2_O in 0.3 mL pyridine at 40 ^°^C, ON. Evaporation of the volatiles with a stream of N_2_ gave a product acetylated at both the 3-position and the imidazole ring.

N-Acetyl-2-(Imidazol-2-ylmethylene)-cholestan-3β-yl acetate. ^1^H NMR (600 MHz, CDCl_3_): 7.241 (1H, d, *J =* 1.5 Hz), 7.018 (1H, d, *J =* 1.5 Hz), 6.823 (1H, s), 5.347 (1H, dd, *J =* 5.4, 11.0 Hz), 3.859 (1H, d, *J =* 13.3 Hz), 2.565 (3H, s), 2.177 (3H, s), 0.891 (3H, d, *J =* 6.4 Hz), 0.860 (3H, d, *J =* 6.6 Hz), 0.855 (3H, d, *J =* 6.6 Hz), 0.716 (3H, s), 0.626 (3H, s).

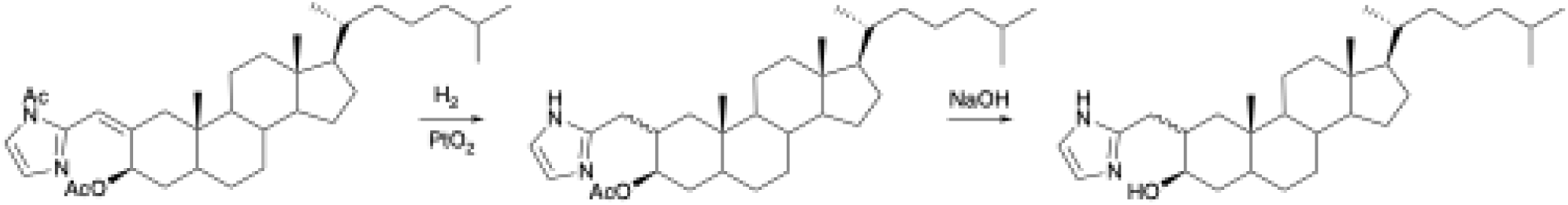

The protected imidazolylmethylene sterol (5 mg) dissolved in 1.25 mL of EtOH was hydrogenated in the presence of 1.5 mg of Adam’s catalyst for 4 hours with stirring, followed by filtration with celite. The product consisted of 63% 2β- and 37% 2α-isomers. Preparative TLC (2% TEA in hexanes/EtOAc 3:2) using a TLC plate pretreated with 2% TEA in hexanes was used to remove the recovered starting material

(20%). The mixture of hydrogenated products (2-3 mg) in 0.2 mL DCM, was saponified with 0.5 mL of 10% NaOH/MeOH with stirring for 1 hr. The reaction was diluted with 7 mL of half-saturated NaHCO_3_ and extracted repeatedly with EtOAc. The organic layers were dried with Na_2_SO_4_ and the product purified by preparative TLC (2% TEA in hexanes/EtOAc 1:2) using a TLC plate pretreated with 2% TEA in hexanes to separate the 2α- and 2β diastereomers.

2β-(1H-Imidazol-2-ylmethyl)-cholestan-3β-ol (2-BMI). ^1^H NMR (600 MHz, CDCl_3_): 6.891 (2H, s), 3.953 (1H, tt, *J =* 5.1,5.8 Hz), 3.307 (1H, dd, *J =* 9.0,15.6 Hz), 2.734 (1H, d, *J =* 15.6 Hz), 2.57-2.52 (1H, m), 0.892 (3H, d, *J =* 6.5 Hz), 0.889 (3H, s), 0.861 (3H, d, *J =* 6.6 Hz), 0.857 (3H, d, *J =* 6.6 Hz), 0.646 (3H, s). ^13^C NMR (201 MHz, CDCl_3_): 150.44, 120.4 (detected by HSQC), 72.84, 56.39, 56.29, 55.25, 45.84, 44.42, 42.66, 40.07, 39.51, 39.05, 36.16, 35.92, 35.79, 35.06, 33.93, 31.95, 30.58, 28.26, 28.23, 28.00, 24.16, 23.83, 22.80, 22.55, 21.30, 18.64, 14.91, 12.14.

### Synthesis of 3β-aminoxy-5β-cholestane (2-BMI)

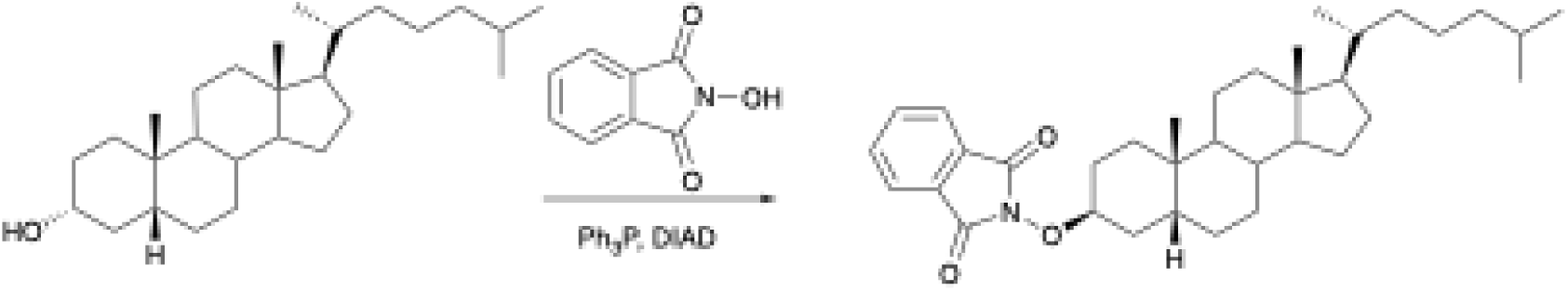

To a solution of 5β-cholestan-3α-ol (17.0 mg, 43.8 μmol) dissolved in 0.5 mL of anhydrous THF under a N_2_ atmosphere was added triphenylphosphine (20.5 mg, 2.0 eq) and *N*-hydroxyphthalimide (12.6 mg, 2.0 eq) and the contents were mixed well to dissolve, followed by the addition of DIAD (15 μL, 2.0 eq). The mixture was allowed to stand at rt for 5.5 hours, then quenched with 5 mL of satd. NaHCO_3_ and extracted thoroughly with hexanes/EtOAc 9:1. The combined organic extracts were filtered through silica which was washed with hexanes/EtOAc 2:1 to ensure complete elution. The product were purified by preparative TLC (hexanes/EtOAc 9:1) to afford the product with inversion of the stereochemistry at C-3 in 90% yield (99% brsm).

3β-Phthalimidooxy-5β-cholestane. ^1^H NMR (600 MHz, CDCl_3_): 7.818 (2H, dd, *J =* 3.1, 5.3 Hz), 7.729 (2H, dd, *J =* 3.1, 5.3 Hz), 4.506 (1H, m), 1.035 (3H, s), 0.899 (3H, d, *J =* 6.5 Hz), 0.863 (3H, d, *J =* 6.6 Hz), 0.858 (3H, d, *J =* 6.6 Hz), 0.659 (3H, s). ^13^C NMR (151 MHz, CDCl_3_): 164.31, 134.29, 129.14, 123.35, 84.02, 56.69, 56.43, 42.76, 40.30, 40.03, 39.52, 36.69, 36.19, 35.77, 35.68, 34.87, 30.09, 29.83, 28.31, 28.00, 26.42, 26.11, 24.22, 23.82, 23.74, 22.78, 22.54, 21.13, 18.69, 12.06.

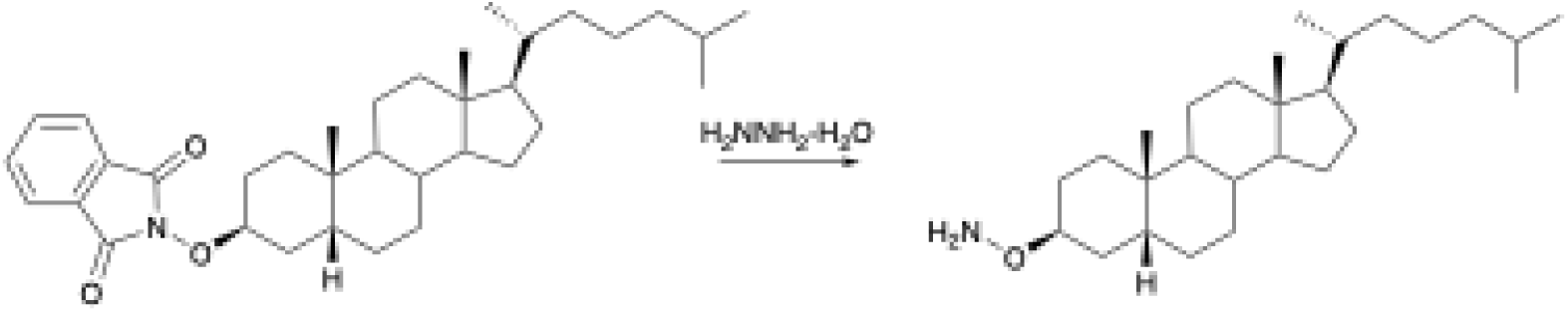

To a solution of the phthalimidooxy intermediate (5.0 mg, 9.4 μmol) in 0.5 mL of anhydrous DCM was added 1 drop (approx. 50 μL) of hydrazine hydrate under a N_2_ atmosphere. The solutions were mixed well and allowed to stand at rt for 2.25 hours. The contents were then filtered through celite and concentrated *in vacuo* at rt to afford the desired product in quantitative yield.

3β-Aminoxy-5β-cholestane (2-BMI). ^1^H NMR (600 MHz, CDCl_3_): 5.165 (2H, br s), 3.790 (1H, m), 1.977 (1H, dt, *J =* 3.5, 12.5 Hz), 1.90-0.95 (m), 0.929 (3H, s), 0.901 (3H, d, *J =* 6.5 Hz), 0.866 (3H, d, *J =* 6.6 Hz), 0.861 (3H, d, *J =* 6.6 Hz), 0.647 (3H, s). ^13^C NMR (151 MHz, CDCl_3_): 79.15, 56.71, 56.44, 42.76, 40.33, 39.95, 39.54, 37.17, 36.21, 35.80, 35.72, 34.97, 30.48, 29.77, 28.33, 28.02, 26.75, 26.22, 24.24, 23.86, 23.85, 23.69, 22.80, 22.56, 21.12, 18.70, 12.07.

### Synthesis of 24-azido-5**α**-chol-5-en-3β-ol

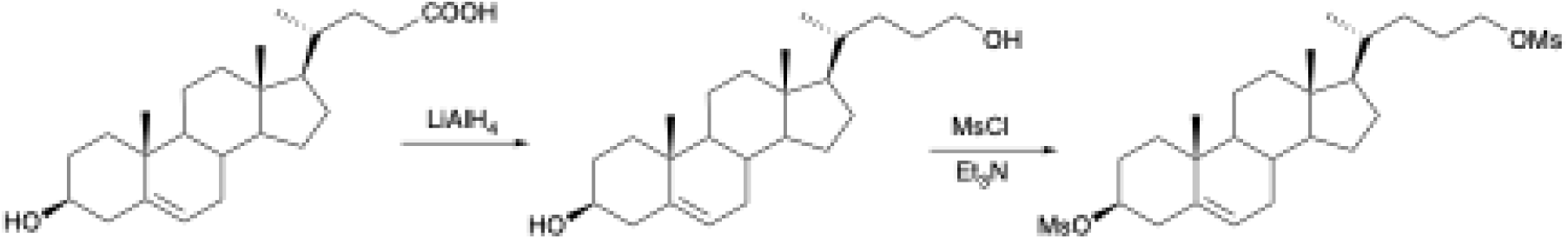

To a solution of 5-cholenic acid (50.3 mg, 0.13 mmol) in 20 mL of anhydrous THF was added solid LAH (62.5 mg, 1.6 mmol, 13 eq.). The mixture was refluxed overnight, then cooled to rt, and excess LAH was destroyed by careful addition of 2 mL of 10% HCl. The mixture was then filtered and concentrated, excess water was removed by azeotrope with benzene. The residue was taken up in EtOAc, filtered through

Na_2_SO_4_, then concentrated to dryness under a stream of N_2_. The crude product was dissolved in 4 mL of anhydrous DCM and 0.1 mL of anhydrous TEA (0.72 mmol), followed by 50 μL of MsCl (0.65 mmol). After 1 hour, complete conversion was observed by TLC and the reaction was quenched with 3.5 mL of 10% HCl and extracted several times with hexanes/EtOAc 2:1. The combined organic extracts were washed with satd. NaHCO_3_, dried over Na_2_SO_4_ and concentrated. Purification by silica gel column chromatography (hexanes/EtOAc 2:1) gave the dimesylate (39.4 mg, 59%).

5α-Chol-5-ene-3β,24-dimesylate ester. ^1^H NMR (600 MHz, CDCl_3_): 5.418 (1H, m), 4.521 (1H, m), 4.24-4.16 (2H, m), 3.005 (3H, s), 3.000 (3H, s), 1.017 (3H, s), 0.939 (3H, d, *J =* 6.5 Hz), 0.680 (3H, s).

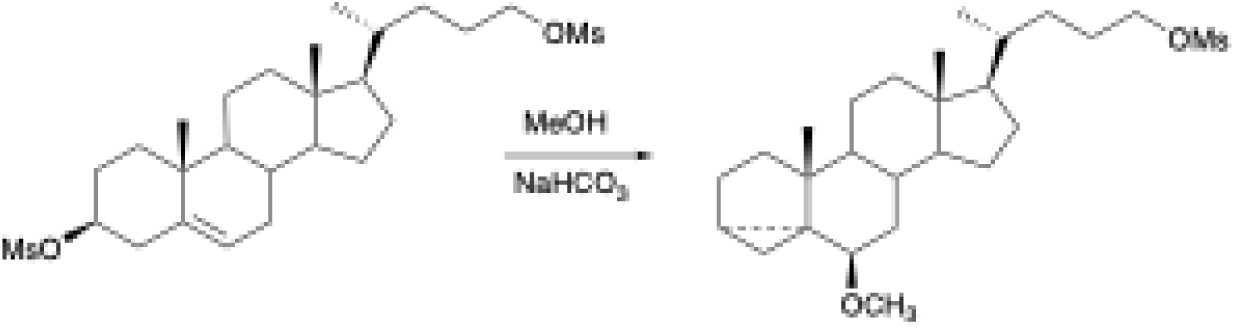

A solution of the dimesylate (19.7 mg, 0.038 mmol) was treated with saturated NaHCO_3_ (350 μL) in methanol (7 mL) at reflux for 1.5 hours. The mixture was diluted with water (40 mL), extracted several times with hexanes/EtOAc 4:1, dried over Na_2_SO_4_ and concentrated. The crude product was purified by preparative TLC (hexanes/EtOAc 4:1) to give 10.2 mg of the i-methyl ether (59%), containing about 9% of the normal methyl ether which was not separated.

24-Methanesulfonoxy-5α-cholane i-methyl ether: ^1^H NMR (600 MHz, CDCl_3_). 4.24-4.16 (2H, m), 3.320 (3H, s), 3.000 (3H, s), 2.768 (1H, t, *J =* 2.6 Hz), 1.019 (3H, s), 0.935 (3H, d, *J =* 6.5 Hz), 0.718 (3H, s).

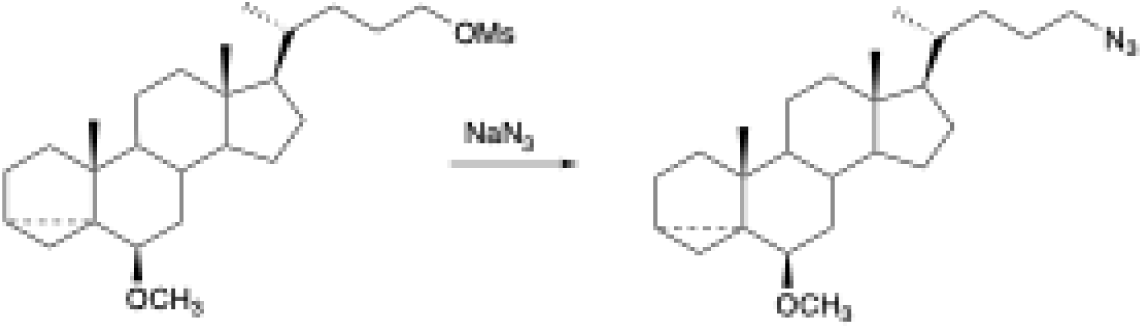

To a solution of the i-methyl mesylate (9.2 mg, 0.02 mmol) in 0.7 mL of anhydrous DMF under N_2_, was added NaN_3_ (7.2 mg, 0.11 mmol). The mixture was stirred at 75 °C for 2 hr, at which point TLC showed complete conversion. The contents were diluted with 7 ml half-saturated NaHCO_3_, extracted thoroughly with hexanes/EtOAc 2:1 and dried over Na_2_SO_4_. Concentration gave 8.0 mg of product (98%) which was used without further purification.

24-Azido-5α-cholane i-methyl ether: ^1^H NMR (600 MHz, CDCl_3_). 3.322 (3H, s), 3.26-3.18 (2H, m), 2.769 (1H, t, *J =* 2.7 Hz), 1.020 (3H, s), 0.933 (3H, d, *J =* 6.5 Hz), 0.719 (3H, s).

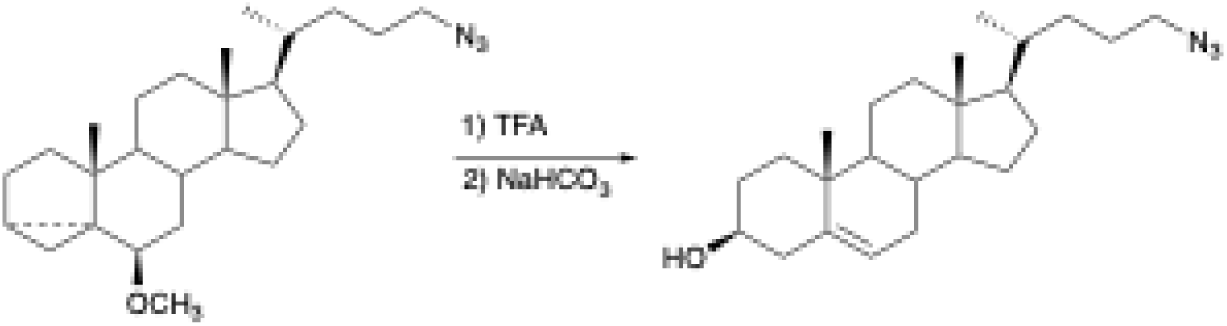

The azido i-methyl ether (7.2 mg, 0.018 mmol) was deprotected in 0.65 mL of DCM containing 1% TFA. When the reaction was found to be complete by TLC, the solution was concentrated to dryness under a stream of N_2_ with gentle heating, and the residue was dissolved in 1 mL of MeOH/DCM 4:1. Three drops of sat. NaHCO_3_ were added and the solution mixed well. The TLC spot at R_f_ = 0.7 was replaced by one at R_f_ = 0.2 (hexanes/EtOAc 4:1) after 20 min. After partitioning between water and hexanes/EtOAc 2:1, the sample was dried with Na_2_SO_4_ and purified by preparative TLC (hexanes/EtOAc 4:1) to give 2.7 mg (38% yield).

24-Azido-5α-chol-5-en-3β-ol: ^1^H NMR (600 MHz, CDCl_3_). 5.350 (1H, m), 3.523 (1H, tt, *J =* 4.5, 11.3 Hz), 3.28-3.18 (2H, m), 1.006 (3H, s), 0.940 (3H, d, *J =* 6.5 Hz), 0.682 (3H, s).

## Notes

### Competing Interest Statement

The authors have declared no competing interest.

